# Sound colless-like balance indices for multifurcating trees

**DOI:** 10.1101/313908

**Authors:** Arnau Mir, Lucía Rotger, Francesc Rosselló

## Abstract

The colless index is one of the most popular and natural balance indices for bifurcating phylogenetic trees, but it makes no sense for multifurcating trees. in this paper we propose a family of *colless-like* balance indices ℭ*_D,f_*, which depend on a dissimilarity *D* and a function *f*: ℕ→ ℝ ⩾_0_, that generalize the colless index to multifurcating phylogenetic trees. We provide two functions *f* such that the most balanced phylogenetic trees according to the corresponding indices ℭ*_D,f_* are exactly the fully symmetric ones. Next, for each one of these two functions *f* and for three popular dissimilarities *D* (the variance, the standard deviation, and the mean deviation from the median), we determine the range of values of ℭ*_D,f_* on the sets of phylogenetic trees with a given number *n* of leaves. We end the paper by assessing the performance of one of these indices on TreeBASE and using it to show that the trees in this database do not seem to follow either the uniform model for multifurcating trees or the *α*-*γ*-model, for any values of *α* and *γ*.

## Introduction

Since the early 1970s, the shapes of phylogenetic trees have been used to test hypothesis about the evolutive forces underlying their assembly [12]. The topological feature of phylogenetic trees most used in this connection is their symmetry, which captures the symmetry of the evolutionary histories described by the phylogenetic trees. The symmetry of a tree is usually measured by means of its *balance* [6, pp. 559–560], the tendency of the children of any given node to have the same number of descendant leaves. Several *balance indices* have been proposed so far to quantify the balance of a phylogenetic tree. The two most popular ones are the *colless index* [4], which only works for bifurcating trees, and the *Sackin index* [17, 20], which can be used on multifurcating trees, but there are many others: see for instance [5, 10, 11, 20] and [6, pp. 562–563].

The *colless index c(T*) of a bifurcating phylogenetic tree *T* is defined as follows: if we call the *balance value* of every internal node *υ* in *T* the absolute value of the difference between the number of descendant leaves of its pair of children, then *c*(*T*) is the sum of the balance values of its internal nodes. in this way, the colless index of a bifurcating tree measures the average balance value of its internal nodes, and therefore it quantifies in a very intuitive way its balance. in particular, *c*(*T*) = 0 if, and only if, *T* is a fully symmetric bifurcating tree with 2*^m^* leaves, for some *m*.

Unfortunately, the colless index can only be used as it stands on bifurcating trees. A natural generalization to multifurcating trees would be to define the balance value of a node as some measure of the spread of the numbers of descendant leaves of its children, like the standard deviation, the mean deviation from the median, or any other dissimilarity applied to these numbers, and then to add up all these balance values. But this definition has a drawback: this sum can be 0 on a non-symmetric multifurcating tree, and hence the resulting index does not capture the symmetry of a tree in a sound way. For an example of this misbehavior, consider the tree depicted in Fig. 1: all children of each one of its nodes have the same number of descendant leaves and therefore the balance value of each node in it would be 0, but the tree is not symmetric. Replacing the number of descendant leaves by the number of descendant nodes, which in a bifurcating tree is simply twice the number of descendant leaves minus 1, does not save the day: again, all children of each node in the tree depicted in Fig. 1 have the same number of descendant nodes.

**Fig 1.**
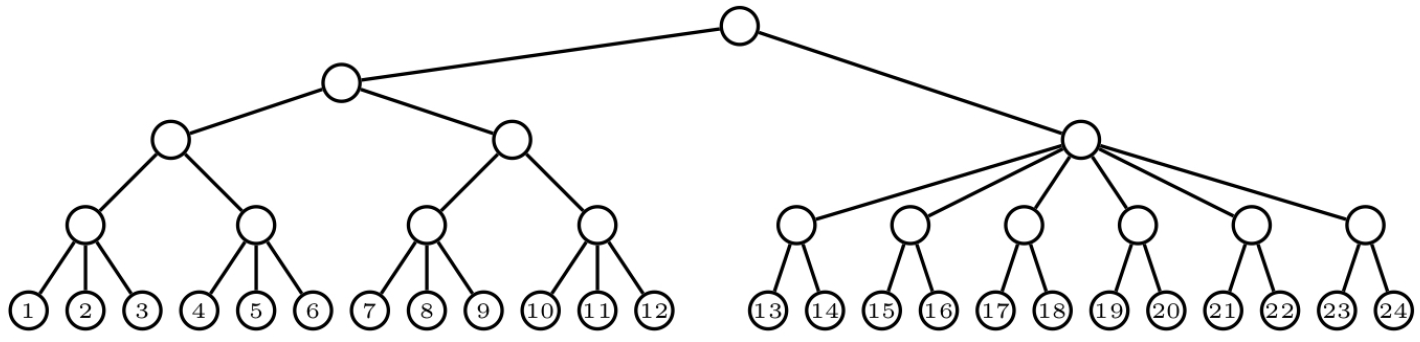
Each node in this asymmetric tree has all its children with the same number of descendant leaves as well as with the same number of descendant nodes.

In this paper we overcome this drawback by taking a suitable function *f*: ℕ → ℝ ⩾_0_ and then replacing in this schema the number of descendant leaves or the number of descendant nodes of a node by the *f-size* of the subtree rooted at the node, defined as the sum of the images under *f* of the degrees of the nodes in the subtree. Then, we define the *balance value* (relative to such a function *f* and a dissimilarity *D*) of an internal node in a phylogenetic tree as the value of *D* applied to the *f*-sizes of the subtrees rooted at the children of the node. Finally, we define the *colless-like index* ℭ*_D,f_* of a phylogenetic tree as the sum of the balance values relative to *f* and *D* of its internal nodes.

The advantage of such a general definition is that there exist functions *f* such that, for every dissimilarity *D*, the resulting index ℭ*_D,f_* satisfies that ℭ*_D,f_* (*T*) = 0 if, and only if, *T* is *fully symmetric*, in the sense that, for every internal node *υ*, the subtrees rooted at the children of *υ* have all the same shape. Two such functions turn out to be *f* (*n*) = ln(*n* + *e*) and *f* (*n*) = *e^n^*.

The different growth pace of these two functions make them to quantify balance in different ways. We show it by finding the trees with largest ℭ*_D,f_* value when *f* is one of these two functions and *D* is the variance, the standard deviation, or the mean deviation from the median. We show that the choice of the dissimilarity *D* does not mean any major difference in the maximally balanced trees relative to ℭ*_D,f_* for a fixed such *f*, but that changing the function *f* implies completely different maximally unbalanced trees.

Finally, we perform some experiments on the TreeBASE phylogenetic database [19]. On the one hand, we compare the behavior of one of our colless-like indices, the one obtained by taking *f* (*n*) = ln(*n* + *e*) and as dissimilarity the mean deviation from the median, MDM, with that of two other balance indices for multifurcating trees: the Sackin index and the total cophenetic index [11]. On the other hand, we use this colless-like index to contrast the goodness of fit of the trees in TreeBASE to the uniform distribution and to the *α*-*γ*-model for multifurcating trees [3].

## Materials

### Notations and conventions

Throughout this paper, by a *tree* we always mean a rooted, finite tree without out-degree 1 nodes. As usual, we understand such a tree as a directed graph, with its arcs pointing away from the root. Given a tree *T*, we shall denote its sets of nodes, of *internal* (that is, non-leaf) nodes, and of arcs by *V* (*T*), *V_int_*(*T*), and *E*(*T*), respectively, and the out-degree of a node *υ* ∈ *V* (*T*) by deg(*υ*). A tree *T* is *bifurcating* when deg(*υ*) = 2 for every *υ* ∈ *V_int_*(*T*). Whenever we want to emphasize the fact that a tree need not be bifurcating, we shall call it *multifurcating*. The *depth* of a node in a tree *T* is the length (i.e., number of arcs) of the directed path from the root to it, and the *depth* of *T* is the largest depth of a leaf in it. We shall always make the abuse of language of saying that two isomorphic trees are equal, and hence we shall always identify any tree with its isomorphism class. We shall denote by 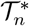 the set of (isomorphism classes of) trees with *n* leaves, and by 𝒯* the union 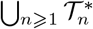.

A *phylogenetic tree* on a (non-empty, finite) set *X* of *labels* is a tree with its leaves bijectively labelled in the set *X*. We shall always identify every leaf in a phylogenetic tree *T* on *X* with its label, and in particular we shall denote its set of leaves by *X*. Two phylogenetic trees *T*_1_*, T*_2_ on *X* are *isomorphic* when there exists an isomorphism of directed graphs between them that preserves the labelling of the leaves. We shall also make always the abuse of language of considering two isomorphic phylogenetic trees as equal. Given a set of labels *X*, we shall denote by 𝒯*_X_* the set of (isomorphism classes of) phylogenetic trees on *X*, and we shall denote by 𝒯_*n*_, for every *n* ⩾ 1, the set 𝒯_{1,2*,⋯,n*}_. Notice that if |*X|* = *n*, then any bijection *X ↔* {1, 2*,…, n*} induces a bijection 𝒯_*X*_ ↔ 𝒯*n*. Moreover, if |*X|* = *n*, there is a forgetful mapping 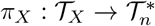 that sends every phylogenetic tree to the corresponding unlabeled tree, which we shall call its *shape*.

No closed formula is known for the numbers 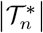 or |𝒯_*n*_|. Felsenstein [6, ch. 3] gives an easy recurrence to compute |𝒯_*n*_| and describes how to obtain such a recurrence for 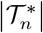; an explicit algorithm to compute the latter is provided in [22]. These numbers |𝒯_*n*_| and 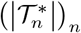 form sequences A000311 and A000669, respectively, in Sloane’s *On-Line Encyclopedia of integer Sequences* [21], where more information about them can be found.

A *comb* is a bifurcating phylogenetic tree with all its internal nodes having a leaf child: see Fig. 2. We shall generically denote every comb in 𝒯_*n*_, as well as their shape in 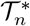, by *K_n_*. A *star* is a phylogenetic tree of depth 1: see Fig. 3. For consistency with later notations, we shall denote the star in 𝒯_*n*_, and its shape in 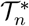, by *FS_n_*.

**Fig 2.**
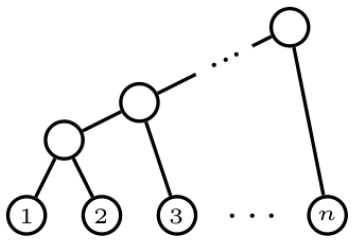
A comb *K_n_* with *n* leaves.

**Fig 3.**
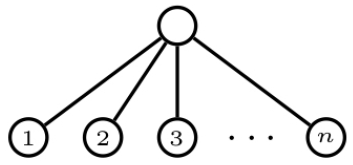
A star *FS _n_* with *n* leaves.

Let *T*_1_*,…, T_k_* be phylogenetic trees on pairwise disjoint sets of labels *X*_1_*,…, X_k_*, respectively. The phylogenetic tree *T*_1_ ⋆ *⋯* ⋆ *T_k_* on *X*_1_ ⋃ *⋯* ⋃ *X_k_* is obtained by adding to the disjoint union of *T*_1_*,…, T_k_* a new node *r* and new arcs from *r* to the root of each *T_i_*. in this way, the trees *T*_1_*,…, T_k_* become the subtrees of *T*_1_ ⋆ *⋯* ⋆ *T_k_* rooted at the children of its root *r*; cf. Fig. 4. A similar construction produces a tree *T*_1_ ⋆ *⋯* ⋆ *T_k_* from a set of (unlabeled) trees *T*_1_*,…, T_k_*.

**Fig 4.**
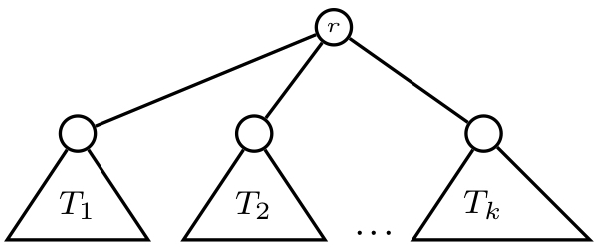
The (phylogenetic) tree *T*_1_ ★ *…* ★ *T_k_*.

Given a node *υ* in a tree *T*, we shall denote by *T_υ_* the subtree of *T* rooted at *υ* and by *κ_υ_* its number of descendant leaves, that is, the number of leaves of *T_υ_*. An internal node *υ* of a tree *T* is *symmetric* when, if *υ*_1_*,…, υ_k_* are its children, the trees *T_υ1_*, *⋯, T_υ_k* are isomorphic. A tree *T* is *fully symmetric* when all its internal nodes are symmetric, and a phylogenetic tree is *fully symmetric* when its shape is so.

Given a number *n* of leaves, there may exist several fully symmetric trees with *n* leaves. For instance, there are three fully symmetric trees with 6 leaves, depicted in Fig. 5. As a matter of fact, every fully symmetric tree with *n* leaves is characterized by an ordered factorization *n*_1_ *⋯ n_k_* of *n*, with *n*_1_*,…, n_k_* ⩾ 2. More specifically, for every *k* ⩾ 1 and (*n*_1_*,…, n_k_*) ∈ ℕ*^k^* with *n*_1_*,…, n_k_* ⩾ 2, let 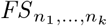 be the tree defined, up to isomorphism, recursively as follows:

• 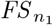 is the star with *n*_1_ leaves.

• If *k* ⩾ 2, 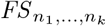 is a tree whose root has *n*_1_ children, and the subtrees at each one of these children are (isomorphic to) 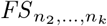.

**Fig 5.**
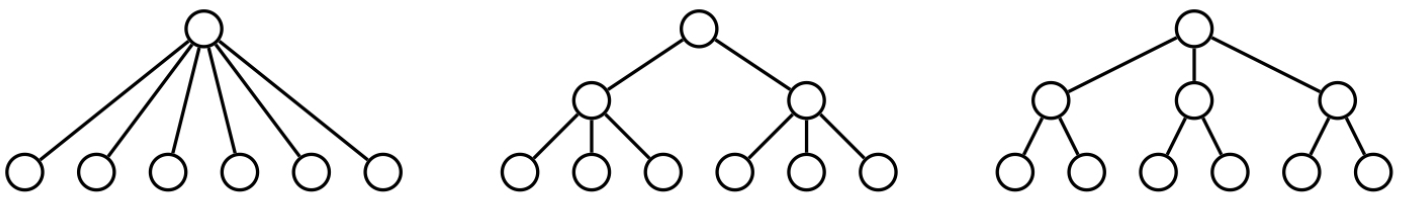
Three fully symmetric trees with 6 leaves: from left to right, *FS* _6_, *FS* _2,3_ and *FS* _3,2_.

Every 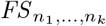 is fully symmetric, and every fully symmetric tree is isomorphic to some 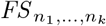. Therefore, for every *n*, the number of fully symmetric trees with *n* leaves is equal to the number *H*(*n*) of ordered factorizations of *n* (sequence A074206 in Sloane’s *On-Line Encyclopedia of integer Sequences* [21]).

### The colless index

The *colless index c*(*T*) of a bifurcating tree *T* with *n* leaves is defined as follows [4]: if, for every *υ* ∈*V_int_*(*T*), we denote by *υ*_1_ and *υ*_2_ its two children and by 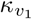 and 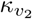 their respective numbers of descendant leaves, then

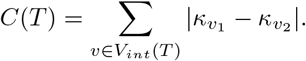

The colless index of a phylogenetic tree is simply defined as the colless index of its shape

It is well-known that the maximum colless index on the set of bifurcating trees with *n* leaves is reached at the comb *K_n_*, and it is

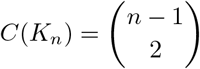

(see, for instance, [16]). As a matter of fact, for every *n* this maximum is only reached at the comb. Since we have not been able to find an explicit reference for this last result in the literature and we shall make use of it later, we provide a proof here.

#### Lemma 1. *For every bifurcating tree T with n leaves, if T ≠ K_n_, then c*(*T*) *< c*(*K_n_*).

*Proof.* Let *T* a bifurcating tree with *n* leaves different from the comb *K_n_*. Let *x* be an internal node of smallest depth in it without any leaf child, and let *T*_1_ ⋆ *T*_2_ and *T*_3_ ⋆ *T*_4_ be the subtrees rooted at its children (see Fig. 6); for every *i* = 1, 2, 3, 4, let *t_i_* be the number of leaves of *T_i_*. Assume, without any loss of generality, that *t*_1_ ⩽*t*_2_ and *t*_1_ + *t*_2_ ⩽ *t*_3_ + *t*_4_. Let then *T′* be the tree obtained by pruning *T*_2_ from *T* and regrafting it to the other arc starting in *x* (see again Fig. 6).

**Fig 6.**
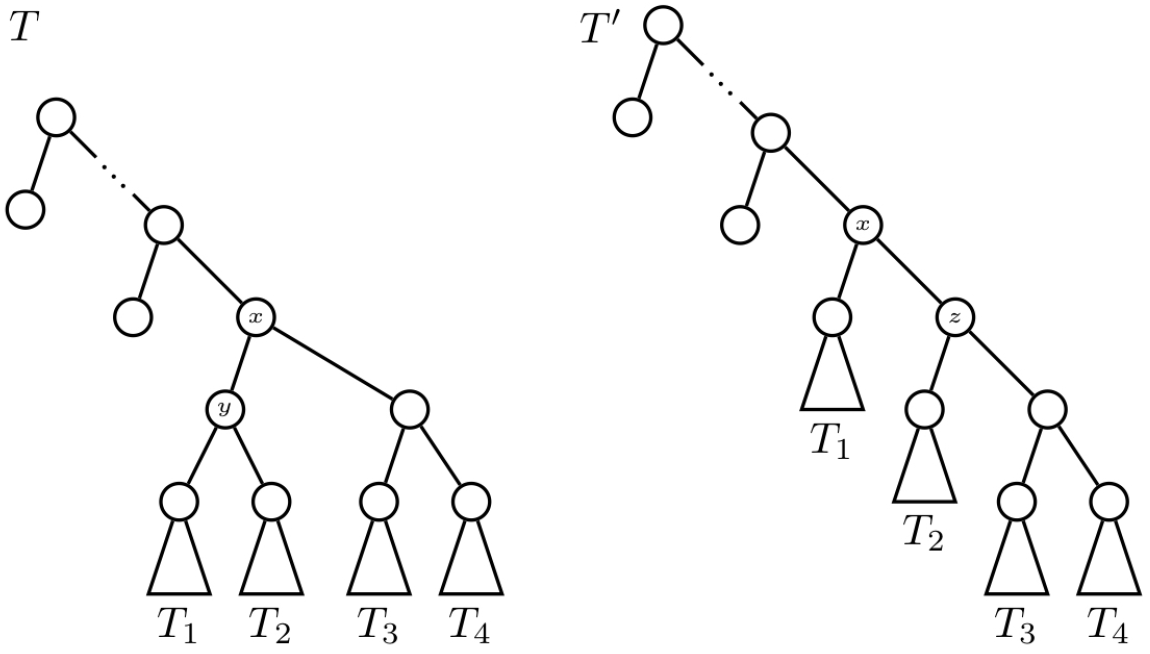
The trees *T* and *T ′* in the proof of Lemma 1.

It turns out that *c*(*T′*) *> c*(*T*). indeed, the only nodes whose children change their numbers of descendant leaves from *T* to *T′* are (cf. Fig. 6): the node *x*; the parent *y* of the roots of *T*_1_ and *T*_2_ in *T*, which is removed in *T′*; and the parent *z* of the root of *T*_2_ in *T′*, which does not exist in *T*. Therefore,

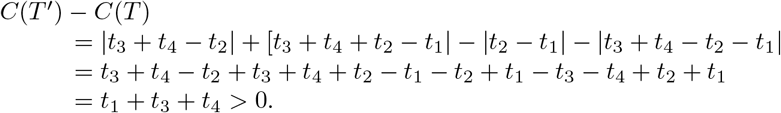

So, this procedure takes a bifurcating tree with *n* leaves *T* ≠ *K_n_* and produces a new bifurcating tree *T′* with the same number *n* of leaves and strictly larger colless index. Since the number of bifurcating trees with *n* leaves is finite, the colless index cannot increase indefinitely, which means that if we iterate this procedure, we must eventually stop at a comb *K_n_*. And since the colless index strictly increases at each iteration, we conclude that if *T ≠ K_n_*, then *c*(*T*) *< c*(*K_n_*).

## Methods

### colless-like indices

Let *f*: ℕ → ℝ_⩾0_ be a function that sends each natural number to a positive real number. The *f-size* of a tree *T* ∈ 𝒯* is defined as

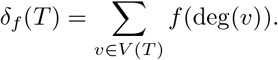

If *T* ∈ *𝒯_X_*, for some set of labels *X*, then *δ_f_* (*T*) is defined as *δ_f_* (*π_X_* (*T*)).

So, *δ_f_* (*T*) is the sum of the degrees of all nodes in *T*, with these degrees weighted by means of the function *f*. Examples of *f*-sizes include:

- The *number of leaves*, *κ*, which is obtained by taking *f* (0) = 1 and *f* (*n*) = 0 if *n >* 0.
- The *order* (the number of nodes), *τ*, which corresponds to *f* (*n*) = 1 for every *n* ∈ ℕ.
- The usual *size* (the number of arcs), *θ*, which corresponds to *f* (*n*) = *n* for every *n*∈ ℕ.

Notice that *δ_f_* satisfies the following recursion:

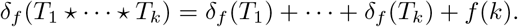

Table 1 in the Supporting File S2 gives the abstract values of *δ_f_* (*T*) for every 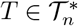 with *n* = 2, 3, 4, 5.

#### Example 2.

*if T is a bifurcating tree with n leaves, and hence with n –* 1 *internal nodes, all of them of out-degree 2, then*

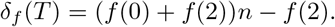

#### Example 3.

*For every fully symmetric tree* 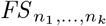,

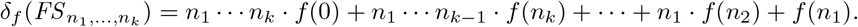

Let now

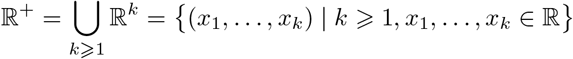

be the set of all non-empty finite-length sequences of real numbers. A *dissimilarity* on ℝ^+^ is any mapping *D*: ℝ^+^ *→* ℝ ⩾_0_ satisfying the following conditions: for every (*x*_1_*,…, x_k_*) ∈ ℝ^+^,

- *D*(*x*_1_*,…, x_k_*) = *D*(*x_σ_*_(1)_*,…, x_σ_*_(*k*)_), for every permutation *σ* ∈ 𝑆_*k*_;
- *D*(*x*_1_*,…, x_k_*) = 0 if, and only if, *x*_1_ = *⋯* = *x_k_*.

The dissimilarities that we shall explicitly use in this paper are the *mean deviation from the median*,

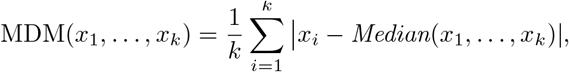

the (*sample*) *variance*,

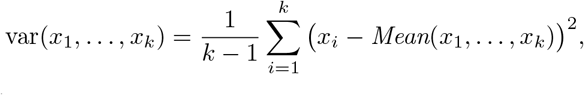

and the (*sample*) *standard deviation*,

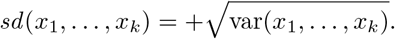

Let *D* be a dissimilarity on ℝ^+^, *f*: ℕ → ℝ_𪟠_ a function, and *δ_f_* the corresponding *f*-size, and let *T* ∈ 𝒯*. For every internal node *υ* in *T*, with children *υ*_1_*,…, υ_k_*, the (*D, f*)*-balance value* of *υ* is

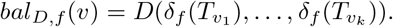

So, *bal_D,f_* (*υ*) measures, through *D*, the spread of the *f*-sizes of the subtrees rooted at the children of *υ*. in particular, *bal_D,f_* (*υ*) = 0 if, and only if, 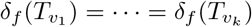.

#### Definition 4.

*Let D be a dissimilarity on* ℝ^+^*and f*:ℕ ℝ⩾_0_ *a function. For every T* ∊*T^*^, its* colless-like index relative to *D* and *f*, ℭ*_D,f_* (*T*)*, is the sum of the* (*D, f*)*-balance values of the internal nodes of T:*

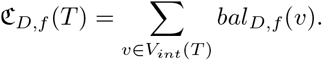

*if T* ∊ *T_X_, for some set of labels X, then* ℭ*_D,f_* (*T*) *is defined as* ℭ*_D,f_* (*π_X_* (*T*)).

#### Example 5.

*if we take D* = MDM *and f the constant mapping 1, so that δ_f_* = *τ, the usual order of a tree, then*

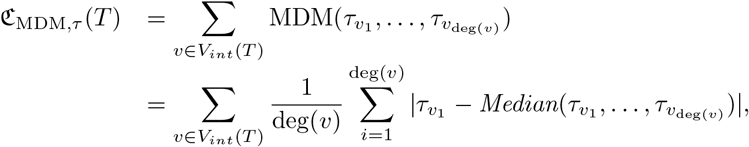

*where, for every v* ∊*V_int_*(*T*)*, v*_1_*,…, v*_deg(*v*)_ *denote its children and τ_v_1__*, *⋯, τ_v_deg(*v*)__ their numbers of descendant nodes.*

Notice that ℭ*_D,f_* gets larger as the *f*-sizes of the subtrees rooted at siblings get more different, and therefore it behaves as a balance index for trees, in the same way as, for instance, the colless index for bifurcating trees: the smaller the value of ℭ*_D,f_* (*T*), the more balanced is *T* relative to the *f*-size *δ_f_*.

It is clear that ℭ*_D,f_* satisfies the following recursion:

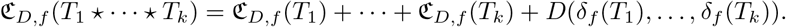

Therefore these colless-like indices are *recursive tree shape statistics* in the sense of [9], relative to the *f*-size *δ_f_*. Table 1 in the Supporting File S2 also gives the abstract values of ℭ*_D,f_* (*T*), for *D* = MDM, var, and *sd*, and for every 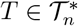 with *n* = 2, 3, 4, 5.

Next result shows that, if we take *D* = MDM or *D* = *sd*, then the restriction of any index ℭ*_D,f_* to bifurcating trees defines, up to a constant factor, the usual colless index.

#### Proposition 6.

*Let T be a bifurcating tree with n leaves and f*: ℕℝ ⩾_0_ *any function. Then*,

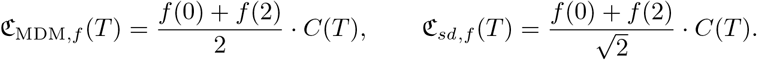

*Proof.* Notice that, for every *x, y* ∊ ℝ, 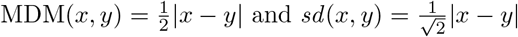 We shall prove the st*√*at<u>e</u>ment for *MDM*; the proof for *sd* is identical, replacing the 2 in the denominator by 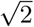. For every internal node *v* in a bifurcating tree *T*, if *v*_1_ and *v*_2_ denote its children,

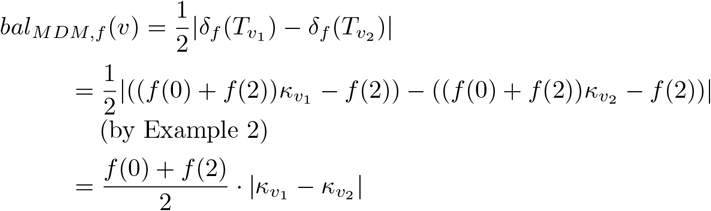

and therefore

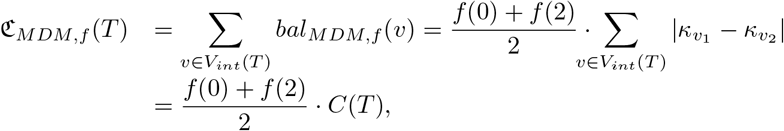

as we claimed. □

If we define the *quadratic colless index* of a bifurcating tree *T* as

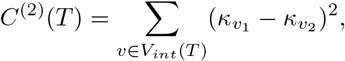

where, for every *v* ∊ *V_int_*(*T*), *v*_1_*, v*_2_ denote its children, then, using that 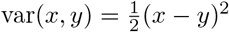, a similar argument proves the following result.

#### Proposition 7.

*Let T be a bifurcating tree with n leaves and f*: ℕ ℝ ⩾0 *any function. Then*,

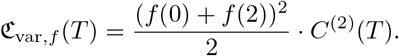
□

As far as the cost of computing colless-like indices goes, we have the following result.

#### Proposition 8.

*if the cost of computing D*(*x*_1_*,…, x_k_*) *is in O*(*k*) *and the cost of computing each f* (*k*) *is at most in O*(*k*)*, then, for every 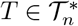, the cost of computing* ℭ*_D,f_* (*T*) *is in O*(*n*).

*Proof.* Assume that every *f* (*k*) is computed in time at most *O*(*k*). For every *k* ⩾2, let *m_k_* the number of internal nodes in *T* of out-degree *k*. Since the sizes *δ_f_* (*v*) are additive, in the sense that if *v* has children *v*_1_*,…, v_k_*, then 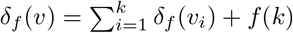, we can compute the whole vector *δ_f_* (*v*) *v*∊_*V* (*T*)_ in time 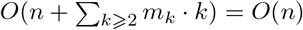 by traversing the tree in post-order.

Assume now that *D*(*x*_1_*,…, x_k_*) can be computed in time *O*(*k*). Then, for every internal node *v* of out-degree *k*, *bal_D,f_* (*v*) = *D*(*δ_f_* (*T_v1_*)*,…, δ_f_* (*T_v_k*)) can be computed in time *O*(*k*), by simply reading the *k* sizes of its children (which are already computed) and applying *D* to them. This shows that the whole vector (*bal _D,f_*(*v*))_*v*∊*V*(*T*)_can be computed again in time 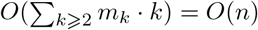. Finally, we compute ℭ_*D,f*_(*T*) by adding the entries of *bal_D,f_* (*v*) *v*∊_*V* (*T*)_, which still can be done in time *O*(*n*).□

The dissimilarities mentioned previously in this subsection can be computed in a number of sums and multiplications that is linear in the length of the input vector, and the specific functions *f* that we shall consider in the next subsection, basically exponentials and logarithms, can be approximated to any desired precision in constant time by using addition and look-up tables [24].

### Sound colless-like indices

It is clear that, for every dissimilarity *D*, for every function *f*: ℕ *→*ℝ ⩾_0_ and for every fully symmetric tree *FS _n_*_1_*,⋯,_nk_*, ℭ*_D,f_* (*FS _n_*_1_*,⋯,_n_k*) = 0, because *bal_D,f_* (*v*) = 0 for every *v*∊*V_int_*(*FS _n_*_1_*,⋯,_nk_*). We shall say that a colless-like index ℭ*_D,f_* is *sound* when the converse implication is true.

#### Definition 9.

*A colless-like index* ℭ*_D,f_* is sound *when, for every T* ∊ *T ∗*, ℭ*_D,f_* (*T*) = 0 *if, and only if, T is fully symmetric.*

In other words, ℭ*_D,f_* is sound when, according to it, the most balanced trees are exactly the fully symmetric trees.

The colless index *c* and its quadratic version *c*^(2)^ are sound for *bifurcating* trees. Unfortunately, neither their direct generalizations ℭ_MDM*,κ*_, ℭ*_sd,κ_* and ℭ*_var,κ_* —where *κ* denotes the number of leaves— nor ℭ_MDM*,τ*_, ℭ*_sd,τ_* and ℭ_var*,τ*_ —where *τ* denotes the number of nodes— or even replacing *τ* by *θ*, the usual size, which is simply *τ* − 1, are sound for multifurcating trees. For example, the tree *T* in Fig. 1 is not fully symmetric, but ℭ_MDM*,κ*_(*T*) = ℭ_var*,κ*_(*T*) = ℭ_*sd,κ*_(*T*) = ℭ_MDM*,τ*_ (*T*) = ℭ_var*,τ*_ (*T*) = ℭ*_sd,τ_* (*T*) = 0.

As a matter of fact, the soundness of ℭ*_D,f_* (*T*) = 0 does not depend on *D*, but only on *f*, as the following lemma shows.

#### Lemma 10.

ℭ*_D,f_ is sound if, and only if, δ_f_* (*T*_1_) ≠ *δ_f_* (*T*_2_) *for every pair of different fully symmetric trees T*_1_*, T*_2_.

*Proof.* As far as the “only if” implication goes, if there exist two different (i.e., non isomorphic) fully symmetric trees *T*_1_*, T*_2_ such that *δ_f_* (*T*_1_) = *δ_f_* (*T*_2_), then the tree *T* = *T*_1_ ⋆ *T*_2_ is not fully symmetric, but

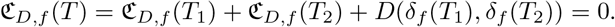

conversely, assume that, for every pair of fully symmetric trees *T*_1_*, T*_2_, if *δ_f_* (*T*_1_) = *δ_f_* (*T*_2_) then *T*_1_ = *T*_2_. We shall prove by complete induction on *n* that if *T* is a tree with *n* leaves such that ℭ*_D,f_* (*T*) = 0, then *T* is fully symmetric. if *T* has only one leaf, it is clearly fully symmetric. Now, assume that *n >* 1 and hence that *T* has depth at least 1. Let *T*_1_*,…, T_k_*, *k* ⩾ 2, be its subtrees rooted at the children of its root, so that *T* = *T*_1_ ⋆ ⋯ ⋆ *T_k_*. Then,

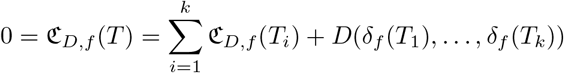
implies, on the one hand, that ℭ*_D,f_* (*T*_1_) = ⋯ = ℭ*_D,f_* (*T_k_*) = 0, and hence, by induction, that *T*_1_*,…, T_k_* are fully symmetric, and, on the other hand, that *D*(*δ_f_* (*T*_1_)*,…, δ_f_* (*T_k_*)) = 0, and hence that *δ_f_* (*T*_1_) = *⋯* = *δ_f_* (*T_k_*), which, by assumption, implies that *T*_1_ = ⋯*⋅*= *T_k_*: in summary, *T* is fully symmetric. □

The following problem now arises:

**Problem.** *To find functions f*: ℕ *→* ℝ_⩾0_ *such that* ℭ*_D,f_ is sound.*

Unfortunately, many natural functions *f* do not define sound colless-like indices, as the following examples show.

#### Example 11.

*if f* (*n*) = *an*^2^ + *bn* + *c, for any a, b, c, then* ℭ*_D,f_ is not sound, because, for example, δ_f_* (*FS* _2,2,2,7_) = *δ_f_* (*FS* _14,4_) = 420*a* + 70*b* + 71*c.*

#### Example 12.

*if f* (*n*) = *n^d^, for any d* ⩾ 0*, then* ℭ*_D,f_ is not sound. indeed, for every d* ⩾ 3 (*the case when d* ⩽ 2 *is a particular case of the last example), take*

- *k* = 2*^d^* + 1 *and l* = 2;
- 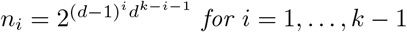
- *n_k_* = 2;
- 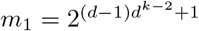
- 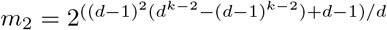*; notice that this exponent is an integer number, because k is odd and therefore d divides* (*d −* 1)*^k^*+ 1.

*Then*

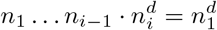

*and hence, on the one hand*,

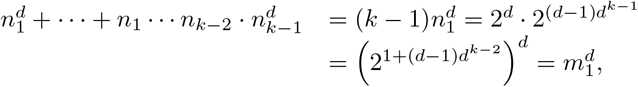

*and, on the other hand*,

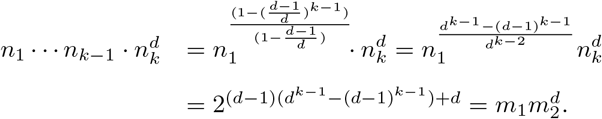

*Therefore*, 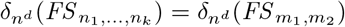

*Of course, for any given d there may exist “smaller” counterexamples: for instance*, 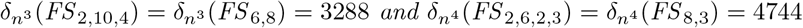.

#### Example 13.

*if f* (*n*) = log_*a*_(*n*) (*for some a >* 1*) when n >* 0*, and f* (0) = 0*, then* ℭ*_D,f_ is not sound: for instance, δ_f_* (*FS* _2,2_) = *δ_f_* (*FS* _8_) = log_*a*_(8)*. in a similar way, if f* (*n*) = log_*a*_(*n* + 1) (*for some a >* 1*), then* ℭ*_D,f_ is not sound, either: for instance, δ_f_* (*FS* _2,3,3_) = *δ_f_* (*FS* _5,7_) = log*a*(196608).

On the positive side, we shall show now two functions that define sound indices. The following lemmas will be useful to prove it.

#### Lemma 14.

*For every k, l* ⩾ 1 *and n*_1_*, n*_2_*,…, n_k_, m*_1_*, m*_2_*,…, m_l_* ⩾ 2*, if* 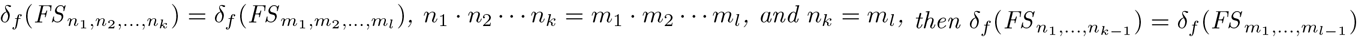.

*Proof.* if *n*_1_ ⋯ *n_k_* = *m*_1_ ⋯ *m_l_* and *n_k_* = *m_l_*, then *n*_1_ ⋯ *n_k−_*_1_ = *m*_1_ ⋯ *m_l−_*_1_. Thus, if, moreover, 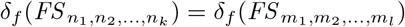, that is,

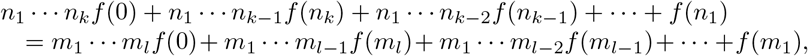

Then

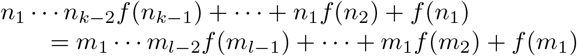

and hence

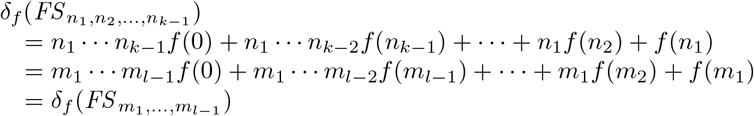

as we claimed. □

#### Lemma 15.

*if n*_1_*,…, n_k_* ⩾ 2*, then*

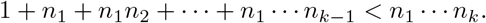

*Proof.* By induction on *k*. if *k* = 1, the statement says that 1 *< n*_1_, which is true by assumption. Assume now that the statement is true for any *n*_1_*,…, n_k_* ⩾ 2, and let *n_k_*_+1_ ⩾ 2. Then,

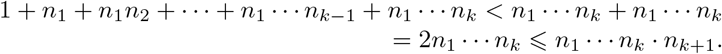
□

#### Proposition 16.

*if f* (*n*) = *e^n^, then* ℭ*_D,f_ is sound.* □

*Proof.* Assume that there exist two non-isomorphic fully symmetric trees 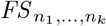 and 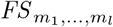 such that

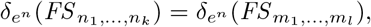

that is, such that

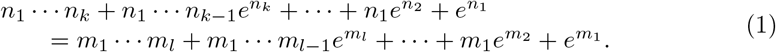

Assume that *l* is the smallest depth of a fully symmetric tree with *e^n^*-size equal to the *e^n^*-size of another fully symmetric tree non-isomorphic to it.

Since *e* is transcendental, equality (1) implies the equality of polynomials in ℤ [*x*]

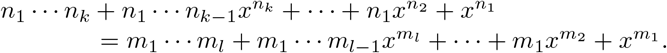

If *l* = 1, the right-hand side polynomial is simply 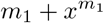 and then the equality of polynomials implies that *k* = 1 and *n*_1_ = *m*_1_, which contradicts the assumption that 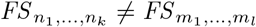. Now assume that *l* ⩾ 2. This equality of polynomials implies the equality of their independent terms: *n*_1_ ⋯ *n_k_* = *m*_1_ ⋯ *m_l_*. On the other hand, the non-zeroth power of *x* with the largest coefficient in the left-hand side polynomial is 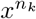 (because all coefficients are non-negative, and, by Lemma 15, *n*_1_ ⋯ *⋅ n_k−_*_1_ alone is larger than the sum *n*_1_ ⋯ *n_k−_*_2_ +⋯+ *n*_1_ + 1 of all other coefficients of non-zeroth powers of *x*) and, by the same reason, the non-zeroth power of *x* with the largest coefficient in the right-hand side polynomial is 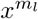. The equality of polynomials implies then that *n_k_* = *m_l_* and hence, by Lemma 14, that 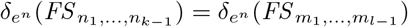, against the assumption on *l*. We reach thus a contradiction that implies that there does not exist any pair of non-isomorphic fully symmetric trees with the same *e^n^*-size. By Lemma 10, this implies that ℭ*_D,e_n* is sound. □

The same argument shows that ℭ*_D,f_* is sound for every exponential function *f* (*n*) = *r^n^* with base *r* a transcendental real number. However, if *r* is not transcendental, then ℭ*_D,r_n* need not be sound. For instance, 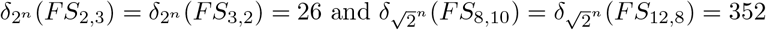.

#### Proposition 17.

*if f* (*n*) = ln(*n* + *e*)*, then* ℭ*_D,f_ is sound.*

*Proof.* The argument is similar to that of the previous proof. Let *f* (*n*) = ln(*n* + *e*) and assume that there exist two non-isomorphic fully symmetric trees 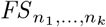 and 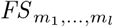 such that 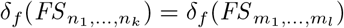, that is, such that

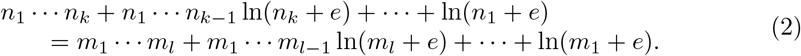

Assume that *l* is the smallest depth of a fully symmetric tree with *f*-size equal to the *f*-size of a fully symmetric tree non-isomorphic to it.

Applying the exponential function to both sides of equality (2), we obtain

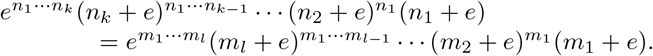

Since *e* is transcendental, this implies the equality of polynomials in ℤ [*x*]

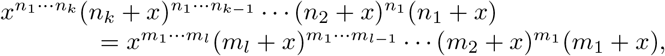

which, since *n*_1_*,…, n_k_, m*_1_*,…, m_l_ ⩾* 2, on its turn implies the equalities

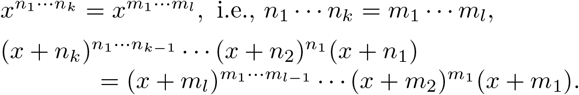

If *l* = 1, the right-hand side polynomial in the second equality is simply *x* + *m*_1_ and then this equality of polynomials implies that *k* = 1 and *n*_1_ = *m*_1_, which contradicts the assumption that 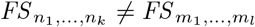. Now assume that *l ⩾* 2. From the first equality we know that *n*_1_ *⋯ n_k_* = *m*_1_ *⋯ m_l_*. Now, the root of the left-hand side polynomial in the second equality with largest multiplicity is *−n_k_* (because, by Lemma 15, *n*_1_ *⋯ n_k−_*_1_ alone is greater than the degree of 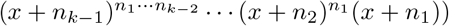 and, similarly, the root of the right-hand side polynomial in the second equality with largest multiplicity is *−m_l_*. Then, the equality of both polynomials implies that *n_k_* = *m_l_* and hence, by Lemma 14, 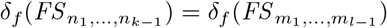, against the assumption on *l*. As in the previous proof, this contradiction implies that ℭ *_D,f_* is sound

The same argument proves that, for every transcendental number *r >* 1, the function *f* (*n*) = log_*r*_(*n* + *r*) defines sound indices ℭ *_D,f_*. However, if *r* is not transcendental, then such a ℭ *_D,f_* need not be sound. For instance,

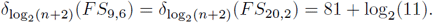

In summary, each one of the functions *f* (*n*) = ln(*n* + *e*) and *f* (*n*) = *e^n^* defines, for every dissimilarity *D*, a colless-like index ℭ *_D,f_* that reaches its minimum value on each 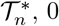, 0, at exactly the fully symmetric trees.

## Results

### Maximally unbalanced trees

The next results give the maximum values of c*_D,f_* on 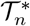 when *D* = MDM, var or *sd* and *f* (*n*) = ln(*n* + *e*) or *f* (*n*) = *e^n^*. These maxima define the range of each ℭ *_D,f_* on 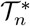, and then, dividing by them, we can define normalized colless-like indices that can be used to compare the balance of trees with different numbers of leaves.

We begin with the function *f* (*n*) = ln(*n* + *e*), which is covered by the following theorem.

#### Theorem 18.

*Let f be a function* ℕ→ ℝ _⩾0_ *such that* 0 *< f* (*k*) *< f* (*k–*1) + *f* (2)*, for every k* ⩾ 3*. Then, for every n* ⩾ 2*, the indices* ℭ*_MDM,f_*, ℭ*_sd,f_ and* ℭ*_var,f_ reach their maximum values on 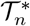 exactly at the comb K_n_. These maximum values are, respectively*,

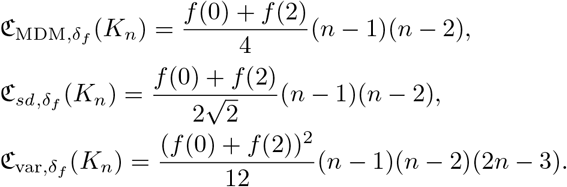
□

The proof of this theorem is very long, and we devote to it the first three sections of the Supporting File S1, one section for each dissimilarity.

It is straightforward to check that the function *f* (*n*) = ln(*n* + *e*) satisfies the hypothesis of Theorem 18 (as to the inequality *f* (*k*) ⩽ *f* (*k–*1) + *f* (2), notice that ln(*k* + *e*) ⩽ ln(*k* + *e –* 1) + ln(2) if, and only if, *k* + *e* ⩽ 2(*k* + *e–*1), and this last inequality holds (strictly) for every *k* ∊ ℕ). Therefore, ℭ_MDM,ln(*n*+*e*)_, ℭ_var,ln(*n*+*e*)_, and ℭ_*sd*,ln(*n*+*e*)_ take their maximum values on 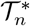 at the comb *K_n_*. in other words, the combs are the most unbalanced trees according to these indices. Table 2 in the Supporting File S2 gives the values of ℭ_MDM,ln(*n*+*e*)_, ℭ_var,ln(*n*+*e*)_, and ℭ_*sd*,ln(*n*+*e*)_ on 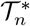 for *n* = 2, 3, 4, 5, and the positions of the different trees in each 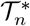 according to the increasing order of the corresponding index.

As far as *f* (*n*) = *e^n^* goes, we have the following result. We have also moved its proof to the Supporting File S1.

#### Theorem 19.

*For every n* ⩾ 2:

(a) *if n ≠* 4*, then both* 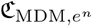 *and* 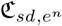 *reach their maximum on* 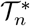 *exactly at the tree FS* _1_ ⋆ *FS _n−_*_1_ (*see Fig. 7), and these maximum values are*

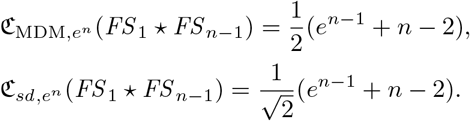
(b) *Both* 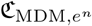 *and* 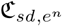 *reach their maximum on* 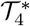 *exactly at the comb K*_4_*, and these maximum values are*

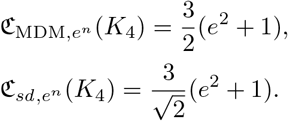
(c) 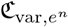 *always reaches its maximum on* 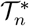 *exactly at the tree FS* _1_ ⋆ *FS _n−_*_1_*, and it is*

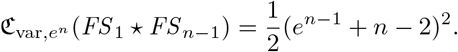

**Fig 7.**
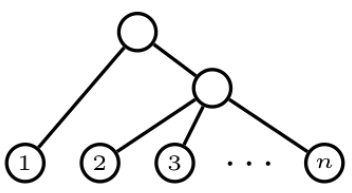
The tree *FS* _1_ ★ *FS _n−_*_1_.

 □

So, according to 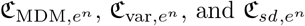, the trees of the form *FS* _1_ ⋆ *FS _n−_*_1_ are the most unbalanced (except for *n* = 4 and *D* = MDM or *sd*, in which case the most unbalanced tree is the comb). Table 2 in the Supporting File S2 also gives the values of these indices on 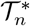, for *n* = 2, 3, 4, 5, and the positions of the different trees in each 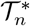 according to the increasing order of the corresponding index.

### The R package “collessLike”

We have written an R package called *collessLike*, which is available at the cRAN (https://cran.r-project.org/web/packages/collessLike/index.html), that computes the colless-like indices and their normalized version, as well as several other balance indices, and simulates the distribution of these indices of 𝒯 *_n_* under the *α*-γ-model [3]. Among others, this package contains functions that:

- compute the following balance indices for multifurcating trees: the Sackin index *S* [17, 20], the total cophenetic index **Φ** [11], and the colles-like index ℭ *_D,f_* for several predefined dissimilarities *D* and functions *f* as well as for any user-defined ones. Our function also computes the normalized versions (obtained by subtracting their minimum value and dividing by their range, so that they take values in [0, 1]) of *S*, Φ and the colless-like indices ℭ *_D,f_* for which we have computed the range in Theorems 18 and 19. Recall from the aforementioned references that, for every *n* ⩾ 2:
  – the range of *S* on 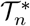 goes from *S*(*FS_n_*) = *n* to 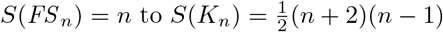
  – the range of Φ on 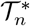 goes from Φ (*FS_n_*) = 0 to 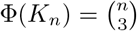
  Therefore, for every T ∊ *𝒯_n_*, the normalized Sackin and total cophenetic index are, respectively,

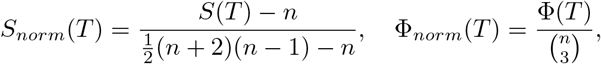

while, for instance, the normalized version of ℭ_MDM,ln(*n*+*e*)_ is

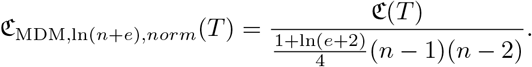
- Given an *n* ⩾ 2, produce a sample of *N* values of a balance index *S*, Φ, or ℭ_*D,f*_ on trees in *𝒯*_n_ generated following an *α*-*γ*-model: the parameters *N, n*, *α*, *γ* (with 0 ⩽ *γ* ⩽ *α* ⩽ 1) can be set by the user. Due to the computational cost of this function, we have stored the values of *S*, Φ, and ℭ_MDM,ln(*n*+*e*)_ (denoted henceforth simply by ℭ) on the samples of *N* = 5000 trees in each *𝒯*_n_(for every *n* = 3, *⋯*, 50 and for every *α*, *γ* ∊ {0, 0.1, 0.2, *⋯*, 0.9, 1 with *γ* ⩽ *α*) generated in the study reported in the next subsection. in this way, if the user is interested in this range of numbers of leaves and this range of parameters, he or she can study the distribution of the corresponding balance index efficiently and quickly.
- Given a tree *T ∊ T_n_*, estimate the percentile 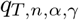 of its balance index *S*, Φ, or ℭ*_D,f_* with respect to the distribution of this index on *𝒯_n_* under some *α*-*γ*-model. if *n*, *α*, *γ* are among those mentioned in the previous item, for the sake of efficiency this function uses the database of computed indices to simulate the distribution of the balance index of *𝒯_n_* under this *α*-*γ*-model.

For instance, the unlabeled tree 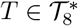 in Fig. 8 is the shape of a phylogenetic tree randomly generated under the *α*-*γ*-model with *α* = 0.7 and *γ* = 0.4 (using set.seed(1000) for reproducibility). The values of its balance indices are given in the figure’s caption.

**Fig 8.**
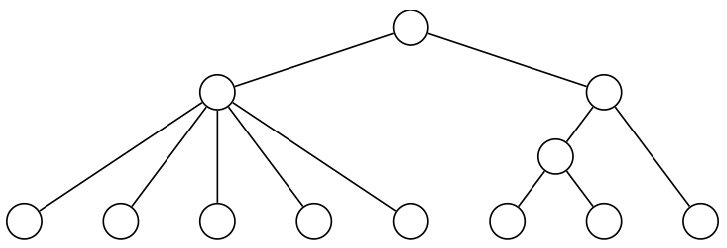
A tree with 8 leaves randomly generated under the *α*-*γ*-model with *α* = 0.7 and *γ* = 0.4. its indices are ℭ (*T*) = 1.746, *S*(*T*) = 18, and Φ(*T*) = 14, and its normalized indices are ℭ _*norm*_(*T*) = 0.06518, *S_norm_*(*T*) = 0.3704, and Φ_*norm*_(*T*) = 0.25.

Fig. 9 displays the estimation of the density function of the balance indices ℭ, *S*, and Φ under the *α*-*γ*-model with *α* = 0.7 and *γ* = 0.4 on *𝒯*_8_, obtained using the 5000 random trees gathered in our database. Moreover, the estimated percentiles of the balance indices of the tree of Fig. 8 are also shown in the figure.

**Fig 9.**
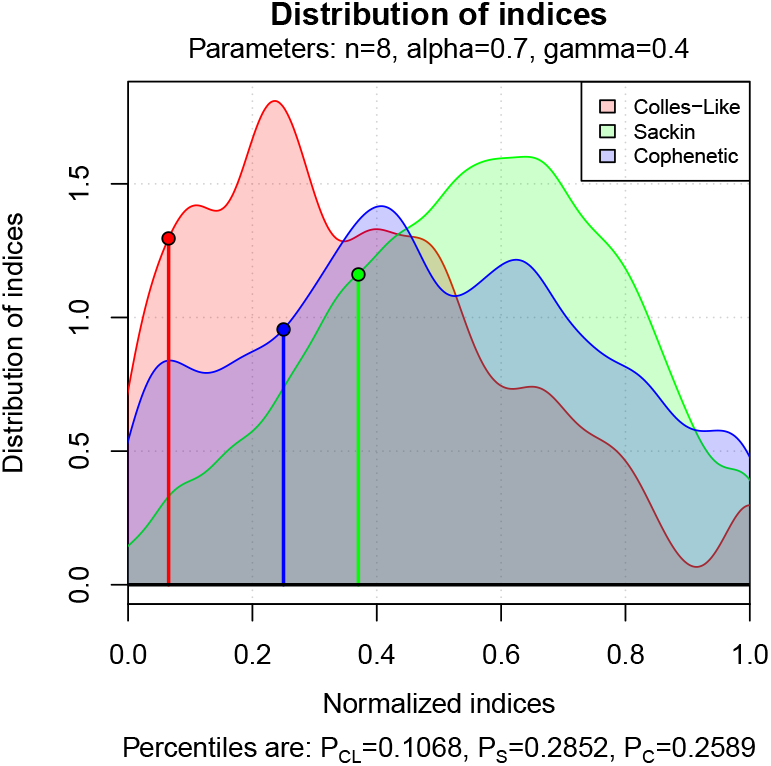
The estimated density function of the distribution of ℭ, *S* and Φ on 𝒯_8_ under the *α*-*γ*-model with *α* = 0.7 and *γ* = 0.4. The percentiles of the tree in Fig. 8 are also represented.

Fig. 10 shows a percentile plot of ℭ, *S*, and Φ under the *α*-*γ*-model for *α* = 0.7 and *γ* = 0.4 on *𝒯*_8_. The percentiles of the tree of Fig. 8 are given by the area to the left of the vertical lines.

**Fig 10.**
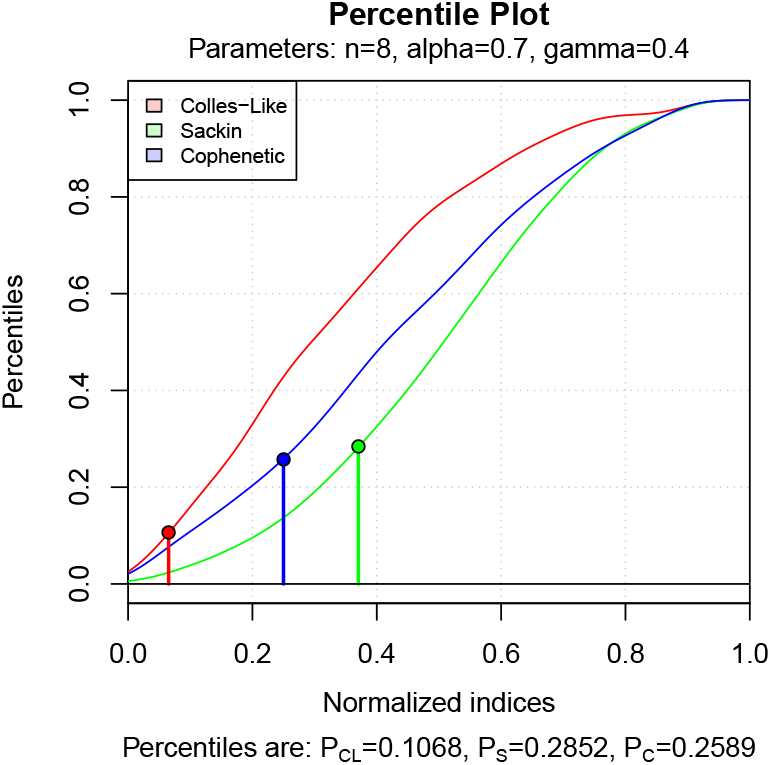
Percentile plot of the distribution of ℭ, *S* and Φ on 𝒯_8_ under the *α*-*γ*-model with *α* = 0.7 and *γ* = 0.4. The percentiles of the tree of Fig. 8 are also highlighted.

Ford’s *α*-model for bifurcating phylogenetic trees [7], which includes as special cases the Yule, or Equal-Rate Markov, model [8, 25] and the uniform, or Proportional to Distinguishable Arrangements, model [2, 14], is on its turn a special case of the *α*-*γ*-model, corresponding to the case *α* = *γ*. So, this package allows also to study this model. For example, the unlabeled tree in Fig. 11 has been generated (with set.seed(1000)) using *n* = 8 and *α* = *γ* = 0.5, which corresponds to the uniform model. The figure also depicts the estimation of the density functions and of the percentile plots of ℭ, *S*, and Φ on *𝒯*_8_ under this model, as well as the percentile values of the tree.

**Fig 11.**
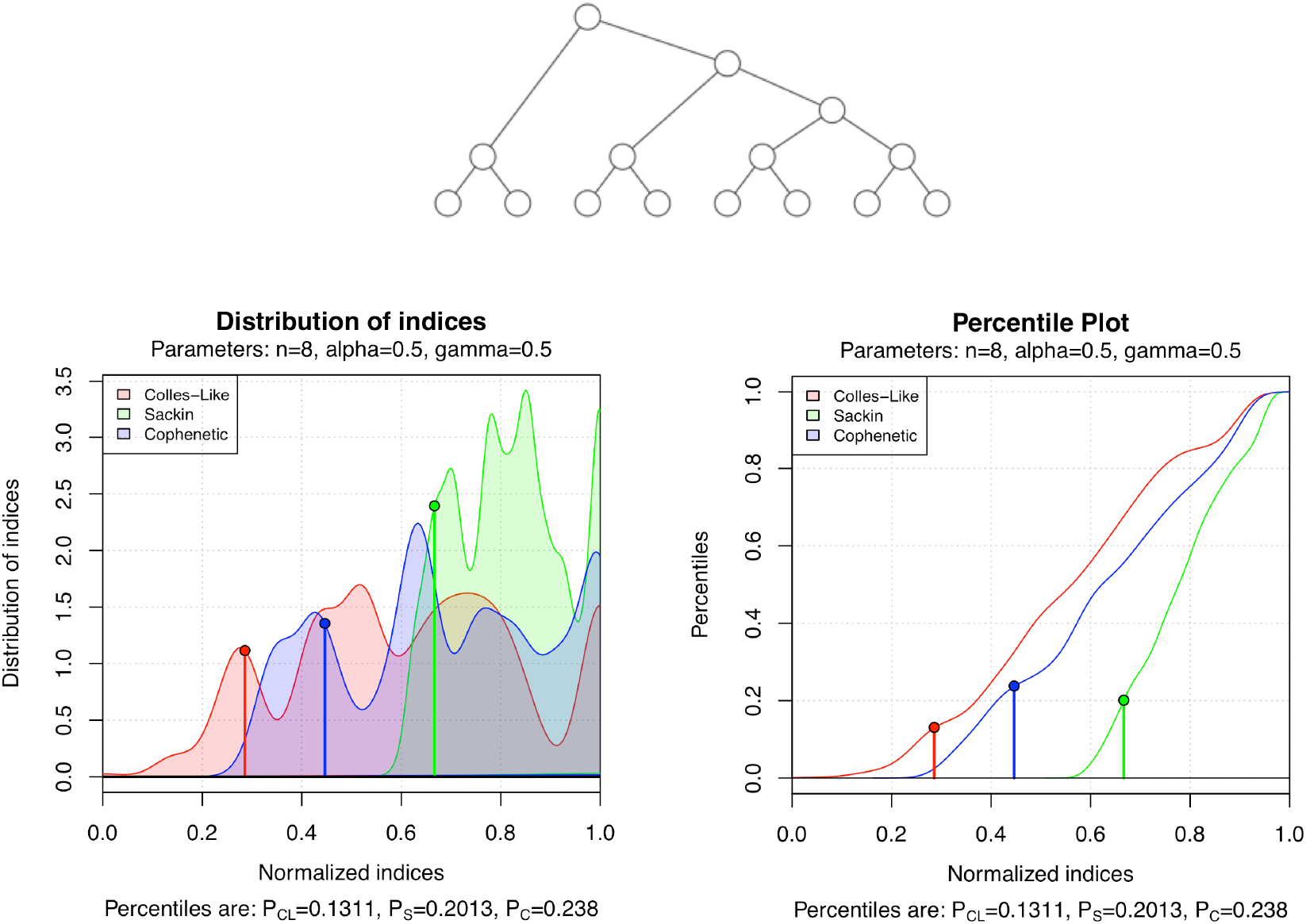
A bifurcating tree randomly generated under the uniform model, the estimated density function of the distribution of the three balance indices on 𝒯_8_ under the uniform model, and their percentile plot.

### Experimental results on TreeBASE

To assess the performance of ℭ_MDM,ln(*n*+e)_, which we abbreviate by ℭ, we downloaded (December 13-14, 2015) all phylogenetic trees in the TreeBASE database [19] using the function search_treebase() of the R package treebase [1]. We obtained 13,008 trees, from which 80 had format problems that prevented R from reading them, so we restricted ourselves to the remaining 12,928 trees. To simplify the language, we shall still refer to this slightly smaller subset of phylogenetic trees as “all trees in TreeBASE”. Only 4,814 among these 12,928 trees in TreeBASE are bifurcating.

Then, for every phylogenetic tree *T* in this set, we have computed its colless-like index ℭ (*T*), its Sackin index *S*(*T*), and its total cophenetic index Φ(*T*). We have then compared the results obtained with ℭ, *S*, and Φ on TreeBASE in the following ways (all analysis have been performed with R [15]).

#### Behavior as functions of the number of leaves

For every number of leaves *n*, we have computed the mean and the variance of ℭ, *S* and Φ on all trees with *n* leaves in TreeBASE. Then, we have computed the regression of these values as a function of *n*.

As far as the means go, the best fits have been:

- *colless-like index*: ℭ̄ ≈ 0.5351 *⋅ n*^1.5848^, with a coefficient of determination of *R*^2^ = 0.9869 and a p-value for the exponent *p <* 2 *⋅* 10^*−*16^.
- *Sackin index*: *S̄ ≈* 1.4512 *⋅ n*^1.4359^, with a coefficient of determination of *R*^2^ = 0.9953 and a p-value for the exponent *p <* 2 *⋅* 10^*−*16^.
- *Total cophenetic index*: Φ̄ *≈* 0.1894 *⋅ n*^2.5478^, with a coefficient of determination of *R*^2^ = 0.9945 and a p-value for the exponent *p <* 2 *⋅* 10^*−*16^.

Fig. 12 depicts these mean values of ℭ (left), *S* (center), and Φ (right) as functions of *n*.

**Fig 12.**
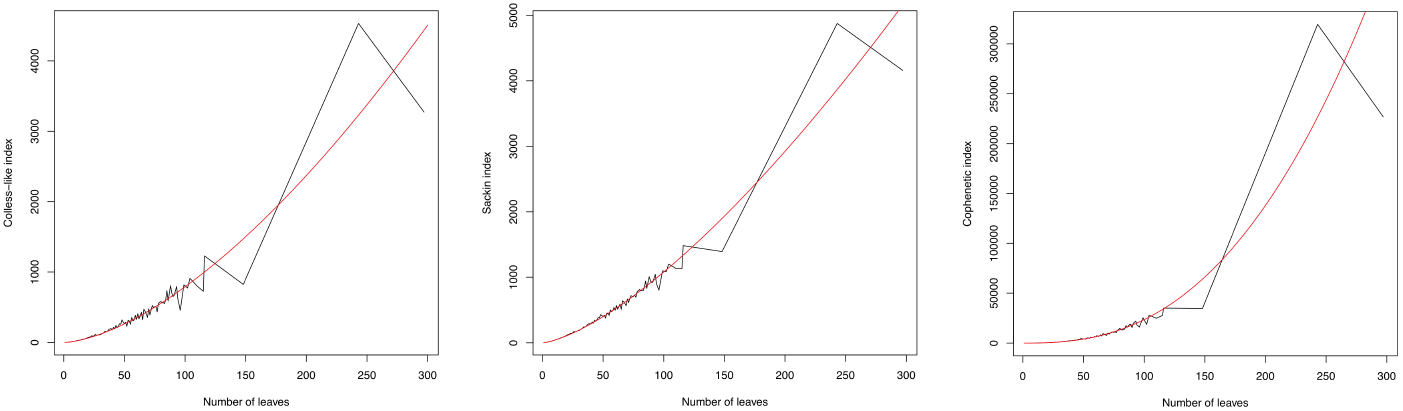
Growth of the mean value of ℭ (left), *S* (center), and Φ (right) in TreeBASE, as functions of the trees’ numbers of leaves *n*.

Thus, *S* and ℭ have similar growth rates, while Φ has a growth rate one order higher in magnitude. This difference vanishes if we normalize the indices by their range, which are *O*(*n*^2^) for ℭ and *S*, and *O*(*n*^3^) for Φ:

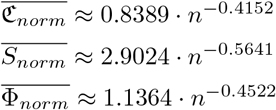

As far as the behavior of the variances goes, the best fits are the following:

- *colless index*: Var(ℭ) *≈* 0.07599 *⋅ n*^3.12831^, with a coefficient of determination of *R*^2^ = 0.962 and a p-value for the exponent *p <* 2 *⋅* 10^*−*16^.
- *Sackin index*: Var(*S*) *≈* 0.03182 *⋅ n*^3.22441^, with a coefficient of determination of *R*^2^ = 0.9575 and a p-value for the exponent *p <* 2 *⋅* 10^*−*16^.
- *Total cophenetic index*: Var(Φ) *≈* 0.0041 *⋅ n*^5.2075^, with a coefficient of determination of *R*^2^ = 0.9812 and a p-value for the exponent *p <* 2 *⋅* 10^*−*16^.

The results are in the same line as before, with the variances of ℭ and *S* having similar growth rates, and the variance of Φ having a growth rate two orders of magnitude higher. This difference vanishes again when we normalize the indices:

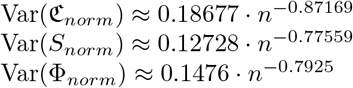

So, in summary, ℭ has, on TreeBASE and relative to the range of values, a slightly larger mean growth rate and a slightly smaller variance growth rate than the other two indices.

#### Numbers of ties

The number of ties (that is, of pairs of different trees with the same index value) of a balance index is an interesting measure of quality, because the smaller its frequency of ties, the bigger its ability to rank the balance of any pair of different trees. Although, in our opinion, this ability need not always be an advantage: for instance, neither Φ nor *S* take the same, minimum, value on all different fully symmetric trees with the same numbers of leaves (for example, *S*(*FS* _6_) = 6 but *S*(*FS* _2,3_) = *S*(*FS* _3,2_) = 12; and Φ(*FS*_6_) = 0, but Φ(*FS* _3,2_) = 3 and Φ(*FS* _2,3_) = 6; cf. Fig. 5), while c applied to any fully symmetric tree is always 0. in this case, we believe that these ties are fair.

Anyway, for every number of leaves *n* and for every one of all three indices under scrutiny, we have computed the numbers of pairs of trees with *n* leaves in TreeBASE having the same value of the corresponding index (in the case of ℭ, up to 16 decimal digits). Fig. 13 plots the frequencies of ties of ℭ, *S* and Φ as functions of *n*. As it can be seen in this graphic, ℭ and Φ have a similar number of ties, and consistently less ties than *S*.

**Fig 13.**
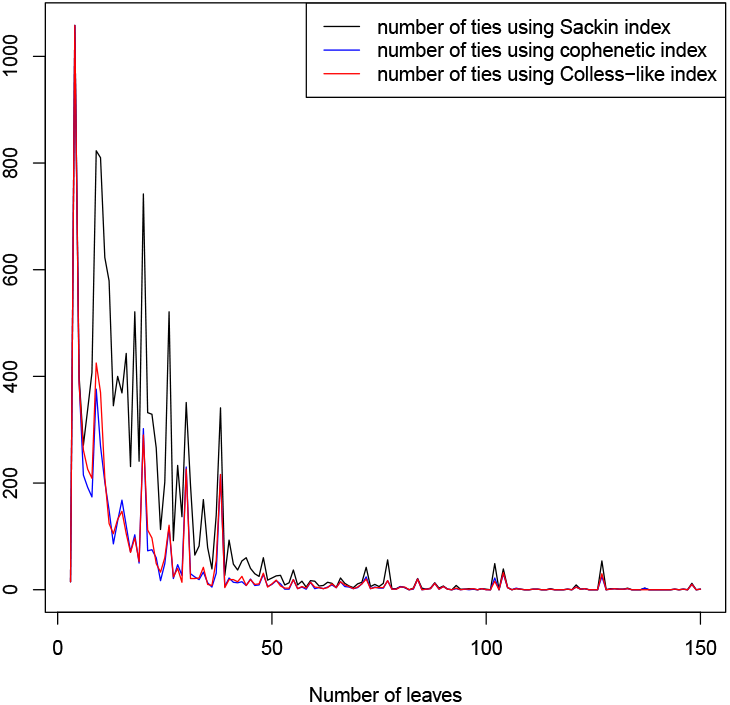
Numbers of ties of ℭ, *S*, and Φ in TreeBASE, as functions of the trees’ numbers of leaves *n*.

#### Spearman’s rank correlation

In order to measure whether all three indices sort the trees according to their balance in the same way or not, we have computed the Spearman’s rank correlation coefficient [18] of the indices on all trees in TreeBASE, as well as grouping them by their number of leaves *n*.

The global Spearman’s rank correlation coefficient of ℭ and *S* is 0.9765, and that of ℭ and Φ is 0.9619. The graphics in Fig. 14 plot these coefficients as functions of *n*. As it can be seen, Spearman’s rank correlation coefficient for ℭ and *S* grows with *n*, approaching to 1, while the coefficient for ℭ and Φ shows a decreasing tendency with *n*.

**Fig 14.**
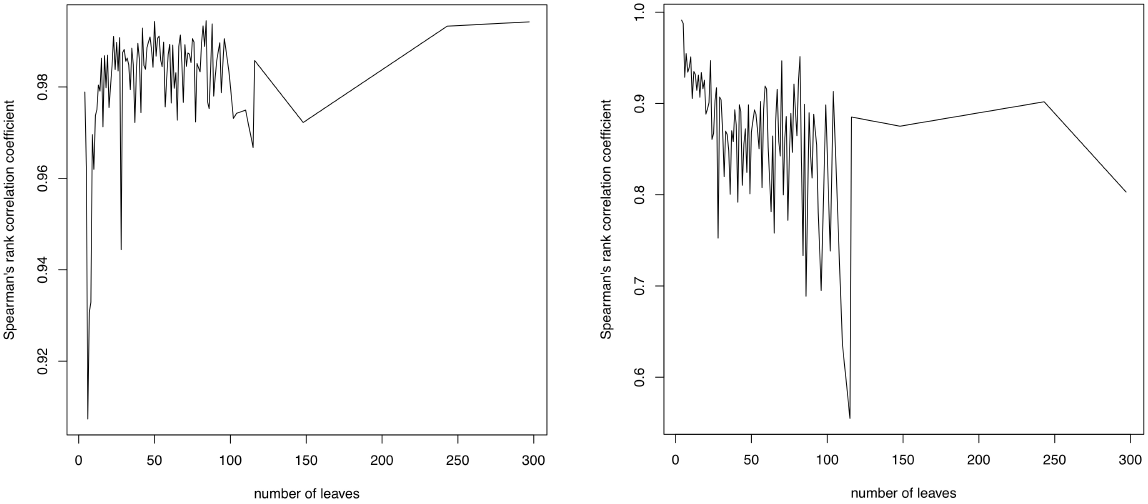
Spearman’s rank correlation coefficient of ℭ and *S* (left) and of ℭ and Φ (right) in TreeBASE, as functions of the trees’ numbers of leaves *n*.

### Does TreeBASE fit the uniform model or the alpha-gamma model?

In this subsection, we test whether the distribution of the colless-like index of the phylogenetic trees in TreeBASE agrees with its theoretical distribution under either the uniform model for multifurcating phylogenetic trees [13] or the *α*-*γ*-model [3] for some parameters *α, γ*. To do it, we use the normalized version ℭ*_norm_* of ℭ, which can be used simultaneously on trees with different numbers of leaves.

To estimate the theoretical distribution of this index under the two aforementioned theoretical models, for every *n* = 3, *⋯*, 50 we have generated, on the one hand, 10,000 random phylogenetic trees in *𝒯_n_* under the uniform model using the algorithm described in [13], and, on the other hand, 5000 random phylogenetic trees in *T_n_* under the *α*-*γ*-model for every pair of parameters (*α, γ*) ∊ {0, 0.1, 0.2, *⋯*, 0.9, 1}^2^ with *γ* ⩽ *α*. We have computed the value of ℭ*_norm_* on all these trees, and we have used the distribution of these values as an estimation of the corresponding theoretical distribution. To test whether the distribution of the normalized colless-like index on TreeBASE (or on some subset of it: see below) fits one of these theoretical distributions, we have performed two non-parametric statistical tests on the observed set of indices of TreeBASE and the corresponding simulated set of indices: Pearson’s chi-squared test and the Kolmogorov-Smirnov test, using bootstrapping techniques in the latter to avoid problems with ties.

As a first approach, we have performed these tests on the whole set of trees in TreeBASE. The p-values obtained in all tests, be it for the uniform model or for any considered pair (*α, γ*), have turned out to be negligible. Then, we conclude confidently that the distribution of the normalized colless-like index on the TreeBASE does not fit either the uniform model or any *α*-*γ*-model when we round *α, γ* to one decimal place. For instance, Fig. 15 displays the distribution of ℭ *_norm_* on TreeBASE and its estimated theoretical distribution under the uniform model. As it can be seen, these distributions are quite different, which confirms the conclusion of the statistical test.

**Fig 15.**
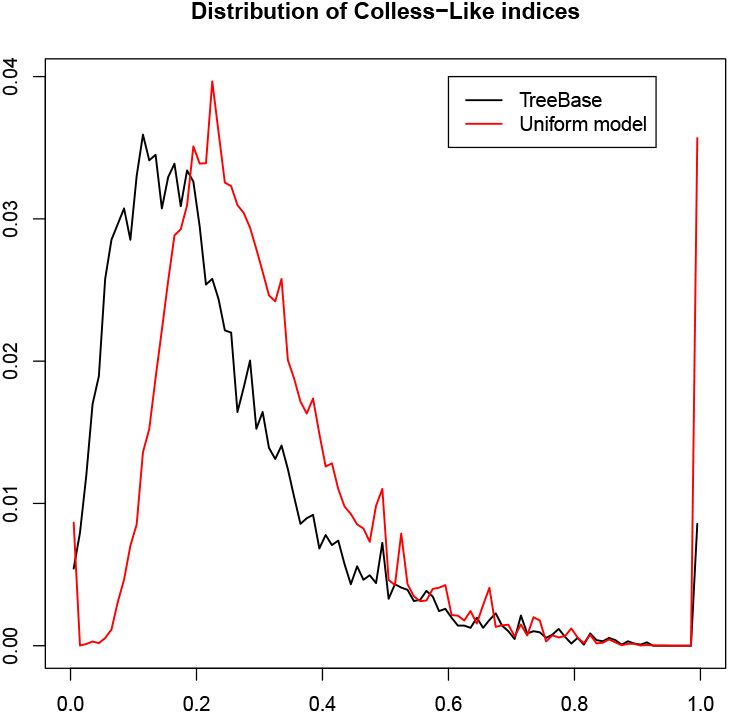
The distribution of ℭ*_norm_* on all trees in TreeBASE (black line) and its estimated theoretical distribution under the uniform model (red line).

Fig. 16 displays the distribution of ℭ *_norm_* for all trees in TreeBASE and its estimated theoretical distribution under the *α*-*γ*-model for the pair of parameters *α, γ* that gave the largest p-values in the goodness of fit tests, which are *α* = 0.7 and *γ* = 0.4. Although graphically both distributions are quite similar, the p-values of the Pearson chi-squared test and of the Kolmogorov-Smirnov test are virtually zero. One might think that the high “peaks” of the theoretical distribution near 0 and 1 could have influenced the outcome of these statistical tests. For this reason, we have repeated them without taking into account these “extreme” values, and the results have been the same.

**Fig 16.**
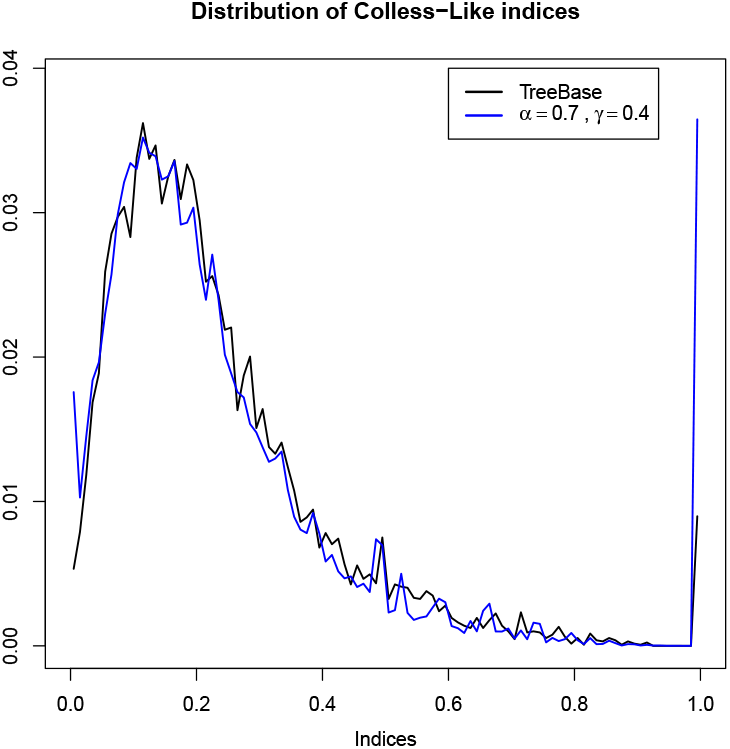
The distribution of ℭ*_norm_* on all trees in TreeBASE (black line) and its estimated theoretical distribution under the *α*-*γ*-model with *α* = 0.7 and *γ* = 0.4 (blue line).

**Fig 17.**
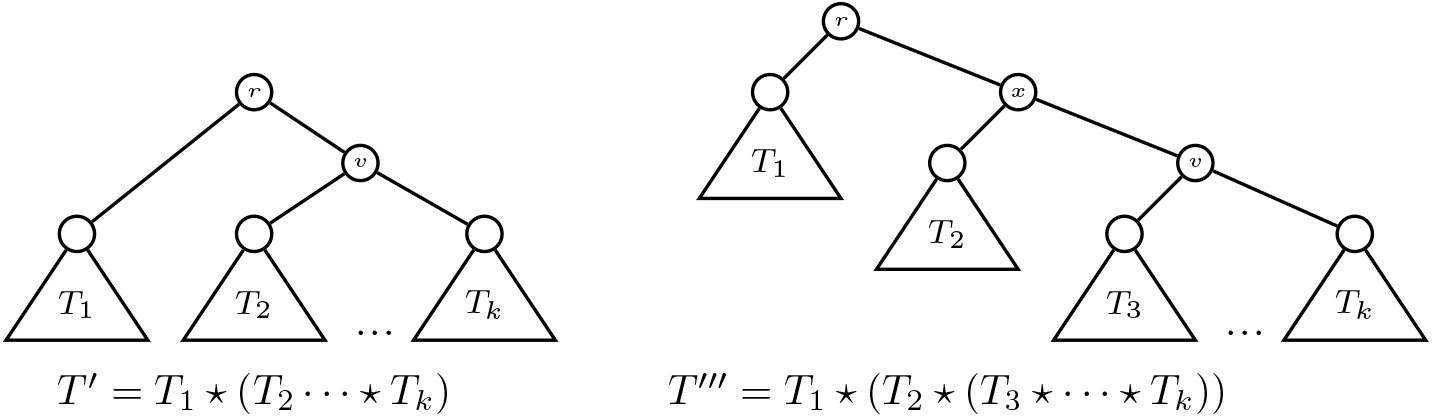
The trees *T′* and *T ^″^* in Lemma 22.

**Fig 18.**
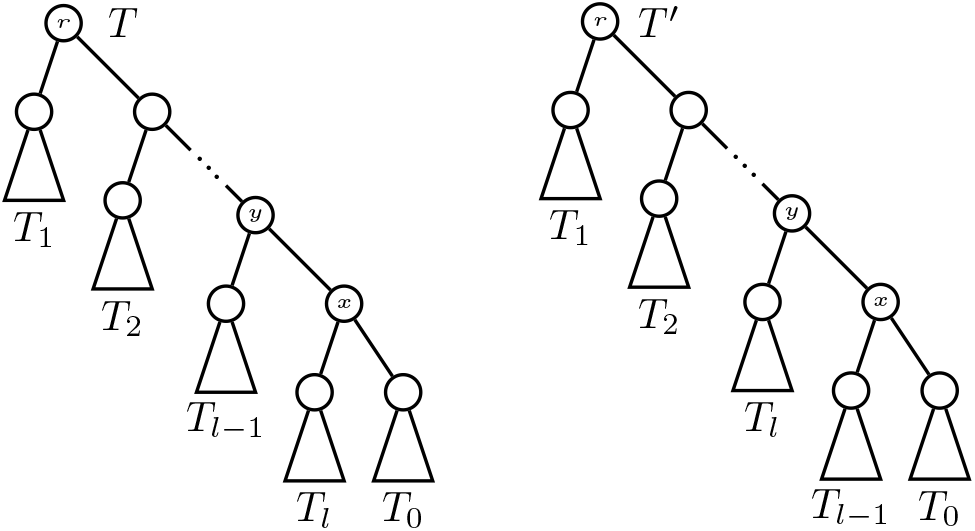
The trees *T* and *T ^′^* in Lemma 23.

**Fig 19.**
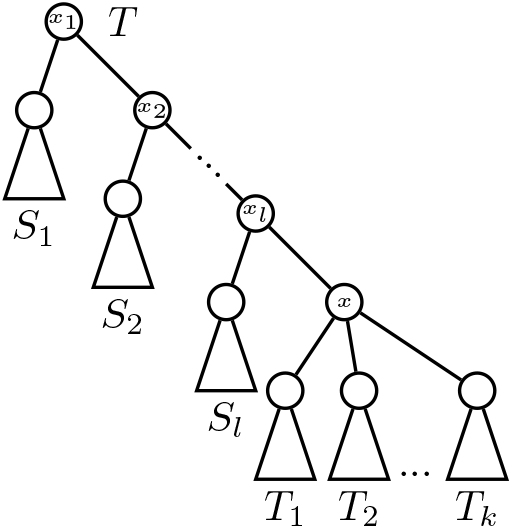
The tree *T* in Lemma 24.

**Fig 20.**
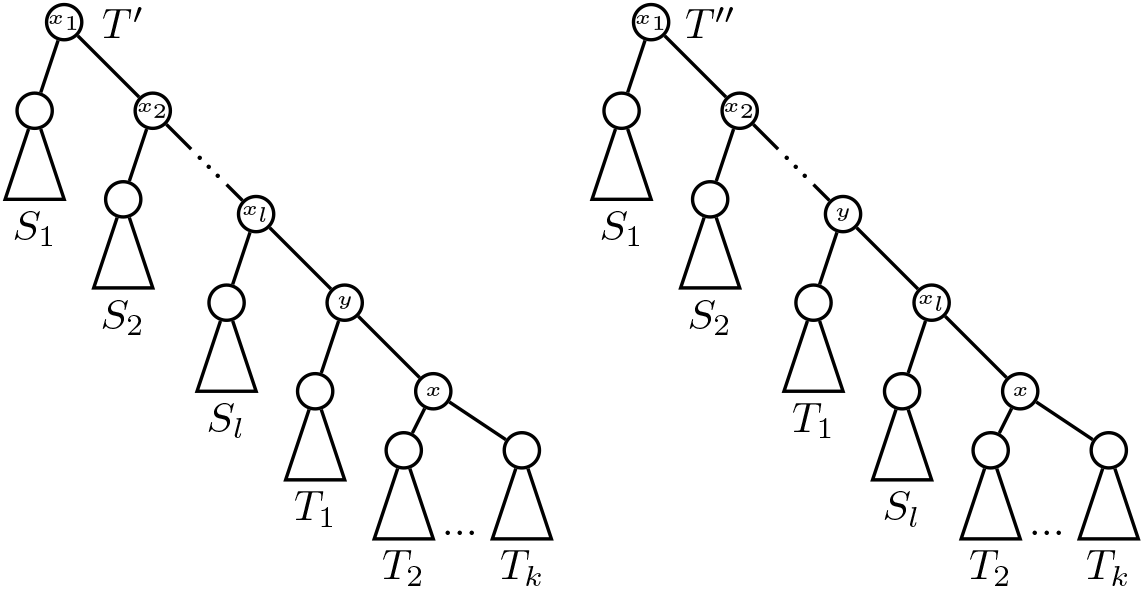
The trees *T ^′^, T ^″^* in Lemma 24.(a).

**Fig 21.**
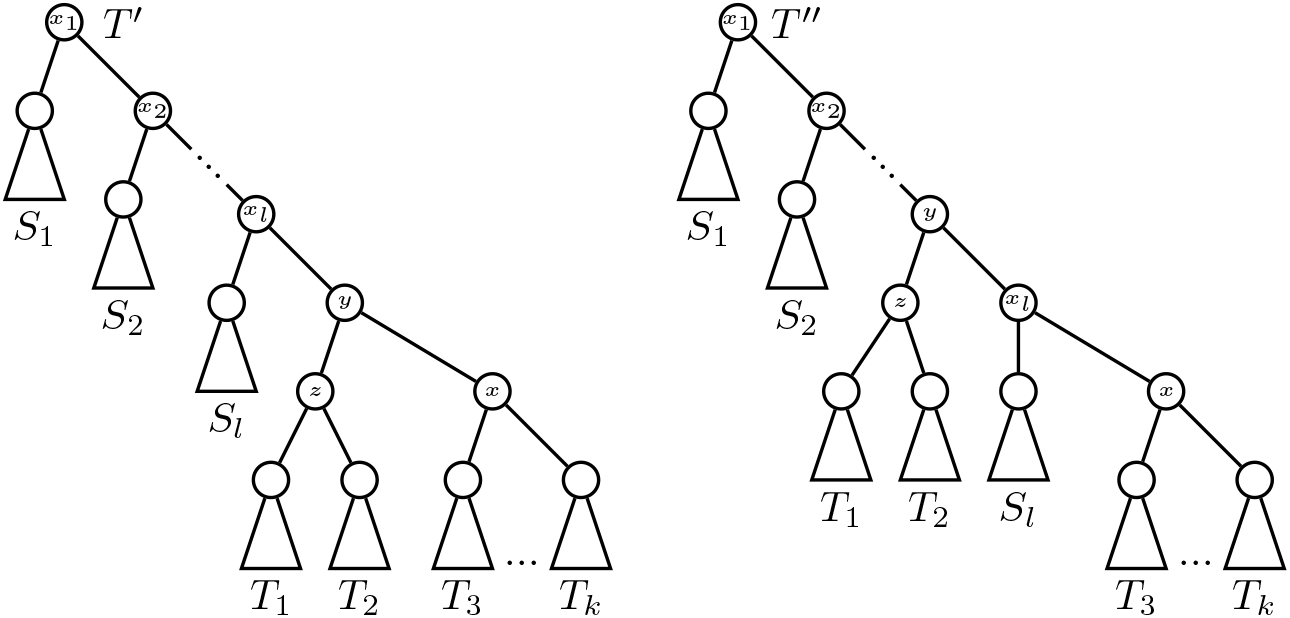
The trees in Lemma 24.(b).

**Fig 22.**
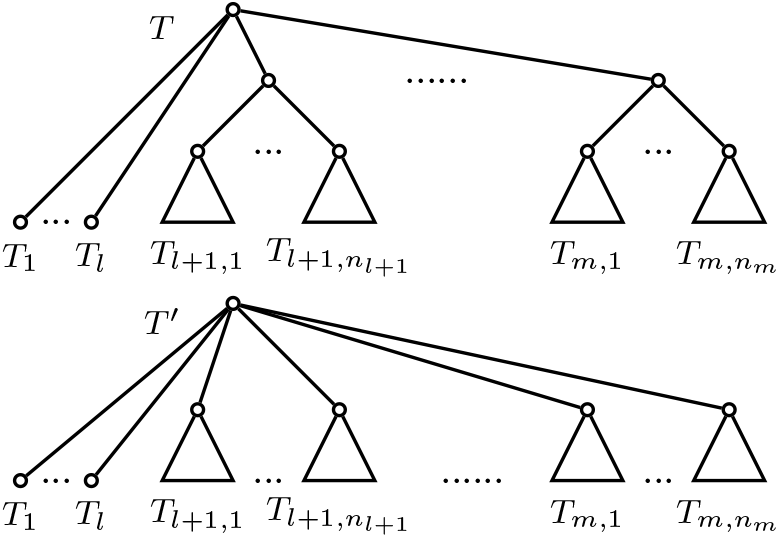
The trees *T* and *T*′ in the proof of Lemma 36.

Since TreeBASE gathers phylogenetic trees of different types and from different sources, we have also considered subsets of it defined by means of attributes. More specifically, besides the whole TreeBASE as explained above, we have also considered the following subsets of it:

- All trees in TreeBASE up to repetitions: we have removed 513 repeated trees (which represent about a 4% of the total).
- All trees with their kind attribute equal to “Species”. This kind attribute can take three values: “Barcode tree”, “Gene Tree” and “Species Tree”.
- All trees with their kind attribute equal to “Species” and their type attribute equal to “consensus”. This type attribute can take two values: “consensus” and “Single”.
- All trees with their kind attribute equal to “Species” and their type attribute equal to “Single”.

We have repeated the study explained above for these four subsets of TreeBASE, comparing the distribution of the normalized colless-like indices of their trees with the estimated theoretical distributions by means of goodness-of-fit tests, and the results have been the same, that is, all p-values have also turned out to be negligible. Our conclusion is, then, that neither the whole TreeBASE nor any of these four subsets of it seem to fit either the uniform model or some *α*-*γ*-model.

## Conclusions

In this paper we have introduced a family of *colless-like* balance indices ℭ *_D,f_*, which depend on a dissimilarity *D* and a function *f*: ℕ → ℝ _⩾0_, that generalize the colless index to multifurcating phylogenetic trees by defining the *balance* of an internal node as the spread, measured through *D*, of the sizes, defined through *f*, of the subtrees rooted at its children. We have proved that every combination of a dissimilarity *D* and a function either *f* (*n*) = ln(*n* + *e*) or *f* (*n*) = *e^n^*, defines a colless-like index that is *sound* in the sense that the maximally balanced trees according to it are exactly the fully symmetric ones. But, the growth of the function *f* used to define the sizes that are compared in the definition of ℭ *_D,f_* determines strongly what the most unbalanced trees are, and hence it has influence on the very notion of “balance” measured by the index.

In our opinion, choosing ln(*n* + *e*) instead of *e^n^* seems a more sensible decision, because, on the one hand, the most unbalanced trees according to the former are the expected ones —the combs— and, on the other hand, we have encountered several hard numeric problems when working with the extremely large figures that appear when using *e^n^*-sizes on trees with internal nodes of high degree. As far a choosing the dissimilarity *D* goes, MDM and *sd* define indices that are proportional to the colless index when applied to bifurcating trees. Among these two options, we recommend to use MDM because it only involves linear operations, and hence it has less numerical precision problems than *sd*, that uses a square root of a sum of squares. This is the reason we have stuck to ℭ _MDM,ln(*n*+*e*)_ in the numerical experiments reported in the Results section.

We want to call the reader’s attention on the problem posed in the “Sound colless-like indices” subsection: to find functions *f* such that ℭ *_D,f_* is sound. Our conjecture is that no function *f*: ℕ *→* ℕ satisfies this property.

## Supporting information

**S1 file: Proofs of Theorems 18 and 19.** The file provides the detailed proofs of Theorems 18 and 19.

**S2 file: Tables.** The file provides two tables, quoted in the main text, with the values of several colless-like indices on 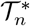 for *n* = 2, 3, 4, 5.

## Competing interests

The authors declare that they have no competing interests.

## Acknowledgements

This research has been partially supported by the *Obra Social la caixa* through the “Programa Pont La caixa per a grups de recerca de la UiB” and by the Spanish Ministry of Economy and competitiveness and European Regional Development Fund through project DPi2015-67082-P (MiNEcO/FEDER). We thank K. Bartoszek for several useful suggestions and A. Saldaña Plomer for making available to us his Java script that generates random phylogenetic trees with a fixed number of leaves with uniform distribution.

## Supplementary files

### Supplementary file S1: Proofs of Theorems 18 and 19

This supplementary document contains the proofs of Theorems 18 (Sections 1–3) and 19 (Sections 4–6) in the main text.

#### 0.1 Proof of the thesis of Theorem 18 for ℭ_MDM,*f*_

Let in this section, and in the next two ones, *f*: ℕ → ℝ_⩾0_ be any mapping such that 0 < *f* (*k*) *< f* (*k−*1) + *f* (2) for every *k* ⩾ 3. Notice that if *f* satisfies this condition then *f* (*k*) *>* 0 not only for every *k* ⩾ 3, but also for *k* = 2, because 0 *< f* (3) *<* 2*f* (2). To simplify the notations, we shall denote in this section *δ_f_* and ℭ_MDM,*f*_ by *δ* and ℭ, respectively, and we shall denote *bal*_MDM,*f*_ on a tree *T* by *bal_T_* or simply by *bal* when it is not necessary to specify the tree.

We split this proof into several lemmas.

##### Lemma 20.

*For every* (*x*_1_*,…, x*_2*n*+1_) ∈ ℝ ^2*n*+1^ *with n* ⩾ 1*, if x_i_ is the median of*

{*x*_1_*,…, x_n_*}*, then*

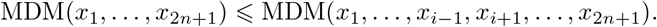

*Moreover, this inequality is strict unless x*_1_ = ⋯ = *x*_2*n*+1_.

*Proof.* After rearranging *x*_1_*,…, x*_2*n*+1_ if necessary, we assume that *x*_1_ ⩽ ⋯ ⩽ *x*_2*n*+1_, in which case their median (i.e., their middle value) is *x_n_*_+1_, and the median of *x*_1_,…*, x_n_, x_n_*_+2_,.…*, x*_2*n*+1_ is *M* = (*x_n_* + *x_n_*_+2_)/2. We want to prove that

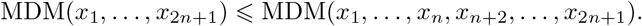

This inequality is true, because

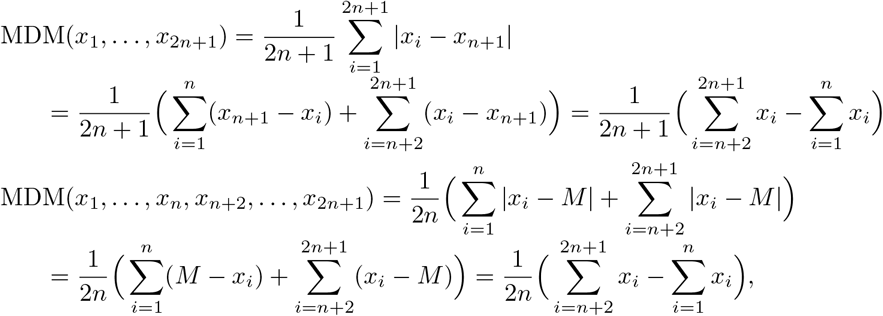

and, clearly,

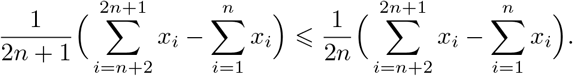

Moreover, the inequality is strict unless 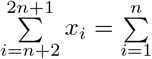, which, under the assumption that *x*_1_ ⩽ ⋯ ⩽ *x*_2*n*+1_, is equivalent to *x*_1_ = ⋯ = *x*_2*n*+1_.

Unfortunately, the thesis of this lemma is false for vectors of numbers of even length. For instance, consider the vector (1, 1, 2, 2): if we remove any single element, its MDM decreases. But, providentially, we can always increase the MDM of an even quantity of real numbers by removing *two* of them.

##### Lemma 21.

*For every* (*x*_1_*,…, x*_2*n*_) ∈ ℝ^2*n*^ with n ⩾ 2*, if x_i_, x_j_, with i < j, are the middle values of* {*x*_1_*,…, x*_2*n*_}*, then*

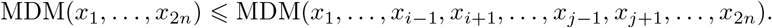

*Moreover, this inequality is strict unless* {*x*_1_*,…, x*_2*n*_} *consists either of* 2*n copies of a single element or of n copies of two different elements.*

*Proof.* After rearranging *x*_1_*,…, x_2n_* if necessary, we assume that *x*_1_ ⩽ ⋯ ⩽ *x*_2*n*_, so that their middle values are *x_n_, x_n_*_+1_, and hence their median is *M* = (*x_n_* + *x_n_*_+1_)/2 and the median of *x*_1_*,…, x_n−_*_1_*, x_n_*_+2_*,…, x_2n_* is *M′* = (*x_n−_*_1_ + *x_n_*_+2_)/2. We want to prove that

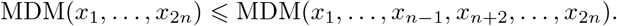

And, indeed,

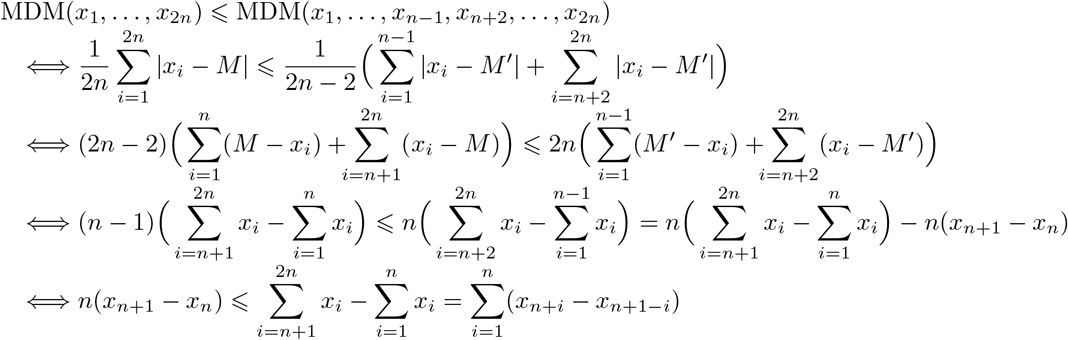

and this last inequality is true because *x_n_*_+1_ *− x_n_* ⩽ *x_n+i_ − x_n+1−i_* for every *i* = 1*,…, n*. Moreover, the inequality is strict unless *x_n_*_+1_ *− x_n_* = *x_n+i_ −*x_n+1*−i*_ for every *i* = 1*,…, n*, that is, unless *x*_1_ = ⋯ = *x_n_* and *x_n_*_+1_ = ⋯ = *x*_2*n*_.

##### Lemma 22.

*Let f be a mapping* ℕ *→* ℝ _⩾ 0_ *such that f* (*k*) *>* 0*, for every k* ⩾ 2*, and let T be a tree of the form T*_1_ ⋆.⋯ ⋆ *T_k_, with k* ⩾ 3 (*see Fig. 4 in the main text).*

(a) *if k is an odd number and if δ*(*T*_1_) *is the median of* {*δ*(*T*_1_)*,…, δ*(*T_k_*)}*, then the tree T′*= *T*_1_ ⋆ (*T*_2_ ⋆ ⋯ ⋆ *T_k_*) (*cf. Fig. 17) satisfies that* ℭ (*T′*) *>* ℭ (*T*).
(b) *if k is an even number and if δ*(*T*_1_)*, δ*(*T*_2_) *are the middle values of* (*δ*(*T*_1_),…, *δ*(*T_k_*)} with *δ*(*T*_1_) ⩽ *δ*(*T*_1_) the the tree *T*″ *T*_1_⋆(*T*_2_ ⋆ (*T*_3_ ⋆ ⋯⋆*T_k_*))(*cf. Fig. 17) satisfies that* ℭ(*T*″) *>* ℭ (*T*).

*Proof.* Let *t_i_* = *δ*(*T_i_*), for every *i* = 1*,…, k*.

As to (a), the only nodes in *T* or *T′* with different *bal* value in both trees are the roots and the new node *v* in *T′*. Therefore,

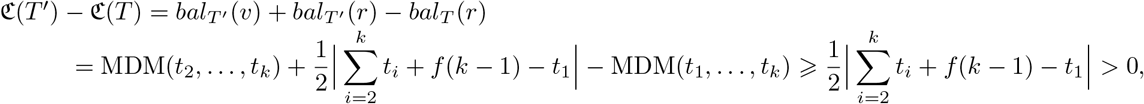

where the first inequality is a consequence of Lemma 20 and the second inequality is strict because, since *t*_1_ is the median of {*t*_1_*,…, t_k_}*and *k* ⩾ 3, there is some *i* ⩾2 such that *t_i_* ⩾ *t*_1_ and hence

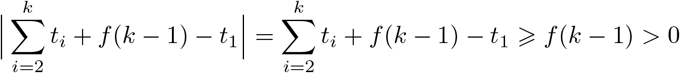

As far as (b) goes, the only nodes in *T* or *T ^″^* with different *bal* value in both trees are the roots and the new nodes *x, v* in *T ^″^*.Therefore,

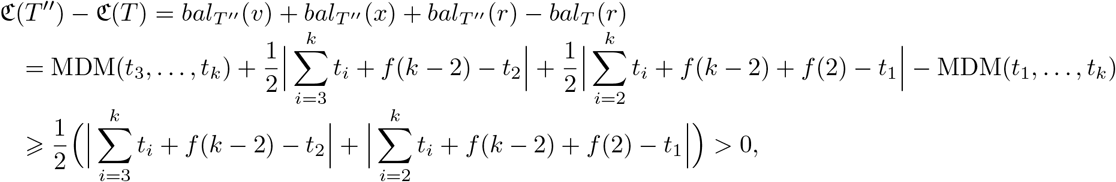

where the first inequality is a consequence of Lemma 21 and the second inequality is strict because, by assumption, *t*_1_ ⩽ *t*_2_ and *k* ⩾ 3, and therefore

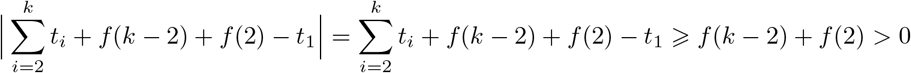

##### Lemma 23.

*Let f be a mapping* ℕ &#211D;⩾0*such that f* (2) *>* 0*. consider the trees T and T ^′^ depicted in Fig. 18, where, in both trees, all nodes in the path from r to x are binary, and T ^′^is obtained from T by simply interchanging the subtrees T_l_ and T_l−_*_1_*. if δ*(*T_l_*) *< δ*(*T_l−_*_1_) *and δ*(*T_l_*) ⩽*δ*(*T*_0_)*, then* &#212D;(*T^′^*) *>* &#212D; (*T*).

*Proof.* Let *t_i_* = *δ*(*T_i_*), for every *i* = 0*,…, l*, so that *t_l_ < t_l−_*_1_ and *t_l_* ⩽ *t*_0_. Since *δ*(*T_y_*) = *δ*(*Ty′*), the only nodes in *T* or *T ^′^* with different *bal* value in both trees are *x* and *y*, and therefore,

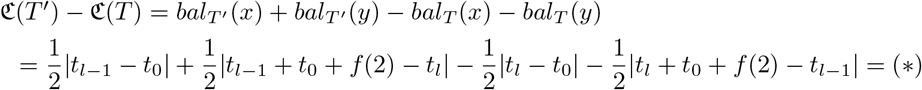

Now we must distinguish two cases:

- If *t_l_ < t_l−_*_1_ ⩽ *t*_0_, then

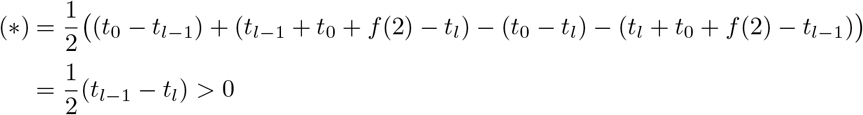
- If *t_l_* ⩽*t*_0_ ⩽*t_l−_*_1_ and *t_l_ < t_l−_*_1_, then

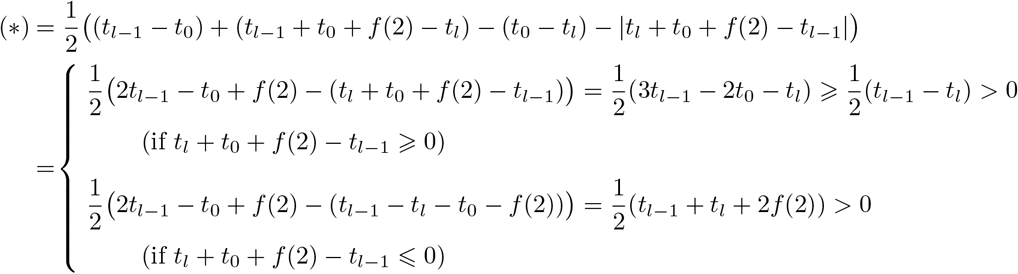

and therefore, in all cases, ℭ (*T ^′^*) *–* ℭ(*T*) *>* 0, as we claimed.

##### Lemma 24.

*Let f be a mapping* ℕ &#211D;⩾0 *such that* 0 *< f* (*k*) *< f* (*k* − 1) + *f* (2)*, for every k* ⩾3*, and let T be the tree depicted in Fig. 19, where l* ⩾1*, x*_1_ *is the root, all nodes in the path from x*_1_ *to x_l_ are binary, and k* ⩾ 3*. Assume moreover that δ*(*S*_1_) ⩽*δ*(*S*_2_) ⩽ *…* ⩽*δ*(*S_l_*).

(a) *Assume that k is odd and that δ*(*T*_1_) *is the median of* {*δ*(*T*_1_)*,…, δ*(*T_k_*)}.

(a.1) *if δ*(*S_l_*) ⩽*δ*(*T*_1_)*, then the tree T^′^ depicted in Fig. 20, obtained by pruning the subtree T*_1_ *and regrafting it in the arc ending in x, satisfies that* ℭ (*T^′^*) *>* ℭ (*T*).
(a.2) *if δ*(*S_l_*) *> δ*(*T*_1_)*, then the tree T ^″^ depicted in Fig. 20, obtained by pruning the subtree T*_1_ *and regrafting it in the arc ending in x_l_, satisfies that* ℭ (*T ^″^*) *>* ℭ (*T*).
(b) *Assume that k is even and that δ*(*T*_1_)*, δ*(*T*_2_) *are the middle values of* {*δ*(*T*_1_)*,…, δ*(*T_k_*)}.

(b.1) *if δ*(*S_l_*) ⩽*δ*(*T*_1_ ⋜ *T*_2_)*, then the tree T ^′^ depicted in Fig. 21, obtained by pruning the subtrees T*_1_ *and T*_2_ *and then inserting T*_1_ ⋜ *T*_2_ *in the arc ending in x, satisfies that* ℭ (*T′*) *>* ℭ (*T*).
(b.2) *if δ*(*S_l_*) *> δ*(*T*_1_ *\* T*_2_)*, then the tree T ^//^ depicted in Fig. 21, obtained by pruning the subtrees T*_1_ *and T*_2_ *and then inserting T*_1_ ⋜ *T*_2_ *in the arc ending in x_l_, satisfies that* ℭ (*T ^″^*) *>* ℭ (*T*).

*Proof.* For every *i* = 1*,…, l*, let *x_i_* denote the parent of the root of *S_i_* in all trees in the statement. Moreover, for every *i* = 1*,…, l*, let *s_i_* = *δ*(*S_i_*) and, for every *i* = 1*,…, k*, let *δ*(*_Ti_*) = *t_i_*, and let 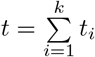. Recall that we are assuming throughout this proof that *s*_1_ ⩽ ⩽*s_l_*.

(*a*) Assume that *k* ⩾3 is odd and that MDM(*t*_1_*,…, t_k_*) ⩽MDM(*t*_2_*,…, t_k_*). As far as assertion (a.1) goes, let us assume that *s_l_* ⩽*t*_1_ and therefore that *s_i_* ⩽*t* for every *i* = 1*,…, l*. in this case, the only nodes in *T* or *T ^′^* with different *bal* value in both trees are *x*_1_*,…, x_l_*, *x* and the new node *y* in *T ^′^.* Then:

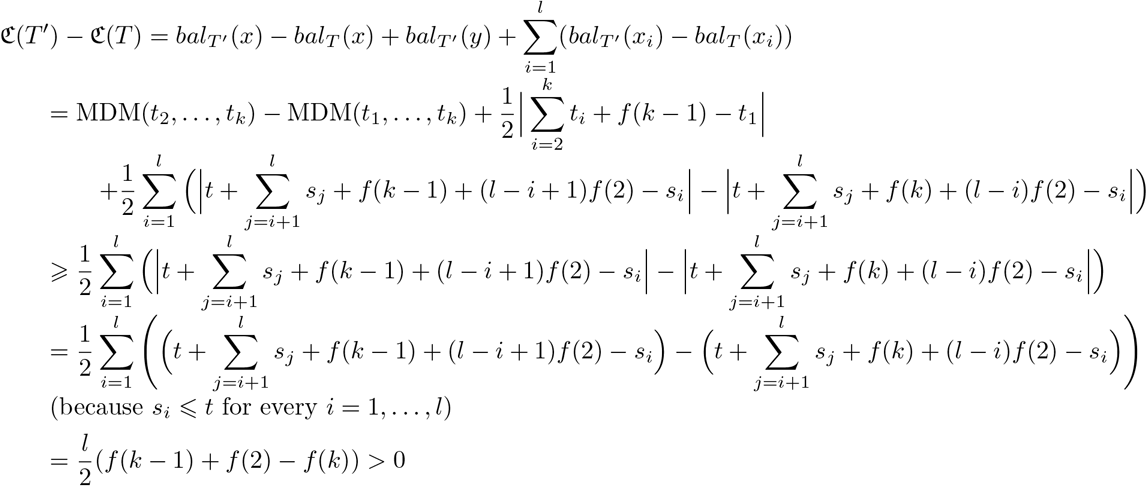 (because *s_i_* ⩽*t* for every *i* = 1*,…, l*) As far as assertion (a.2) goes, let us assume that *s_l_ > t*_1_. Again, the only nodes in *T* or *T ^″^* with different *bal* value in both trees are *x*_1_*,…, x_l_*, *x* and the new node *y*. Therefore:

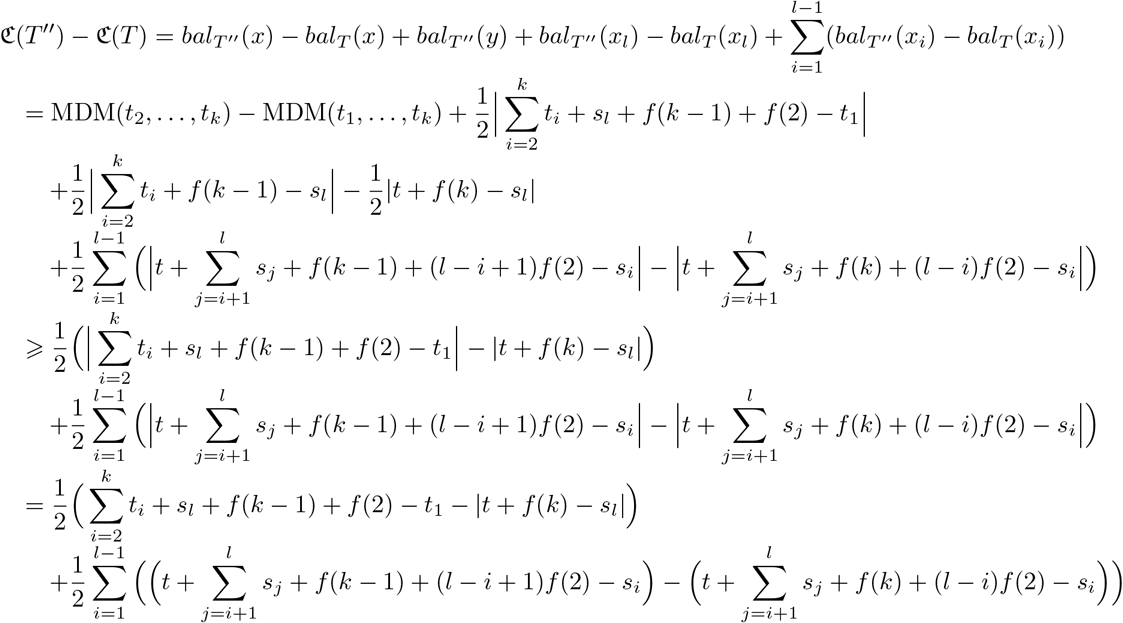 (because *s_i_* ⩽*s_l_*, for every *i* = 1,…*, l*, and *s_l_ > t*_1_)

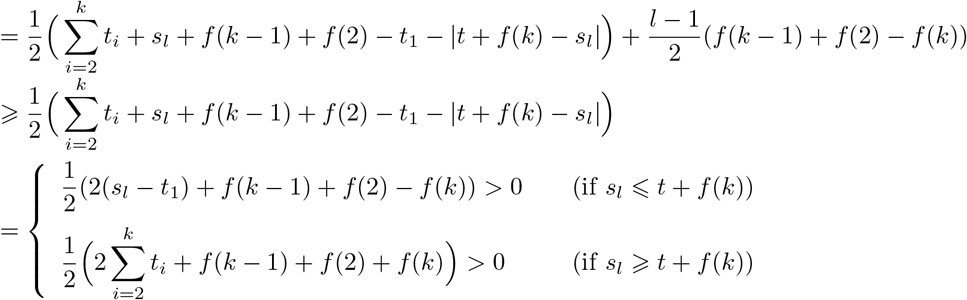
(*b*) Assume now that *k* ⩾3 is even, and hence *k* ⩾4, and that MDM(*t*_1_*,…, t_k_*) ⩽ MDM(*t*_3_,…, *t_k_*). As far as assertion (b.1) goes, let us assume that *s_l_* ⩽ *t*_1_ + *t*_2_ + *f* (2) ⩽ *t* + *f* (2). The only nodes in *T* or *T′* with different *bal* value in both trees are *x*_1_,…, *x_l_*, *x* and the new nodes *y* and *z* in *T*′. Therefore:

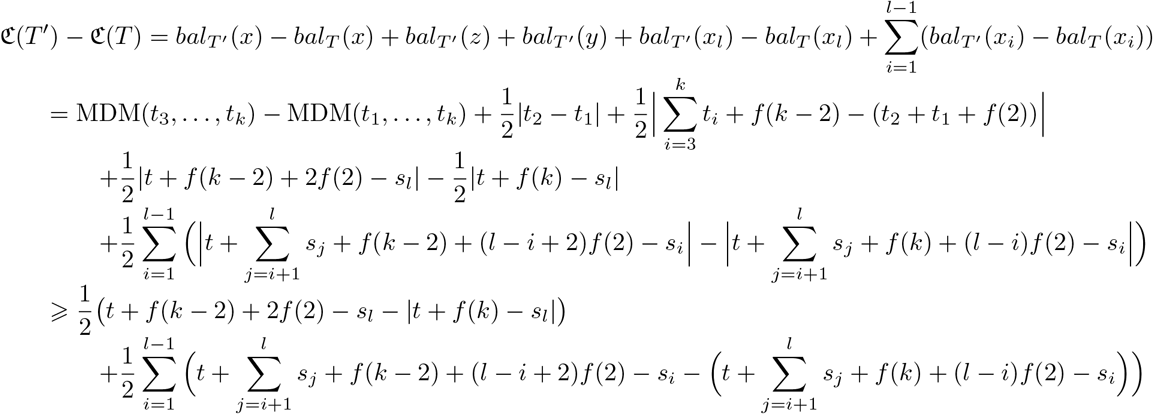

(because *s_l_* ⩽ *t* + *f* (2) and *s_i_* ⩽ *s_l_*, for every *i* = 1,…, *l −* 1)

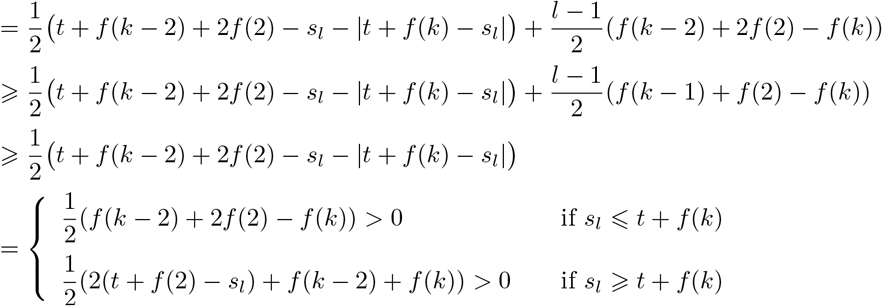 As far as assertion (b.2) goes, let us assume now that *s_l_ > t*_1_ + *t*_2_ + *f* (2). Again, the only nodes in *T* or *T″* with different *bal* value in both trees are *x*_1_,…, *x_l_*, *x* and the new nodes *y*, *z*. Therefore:

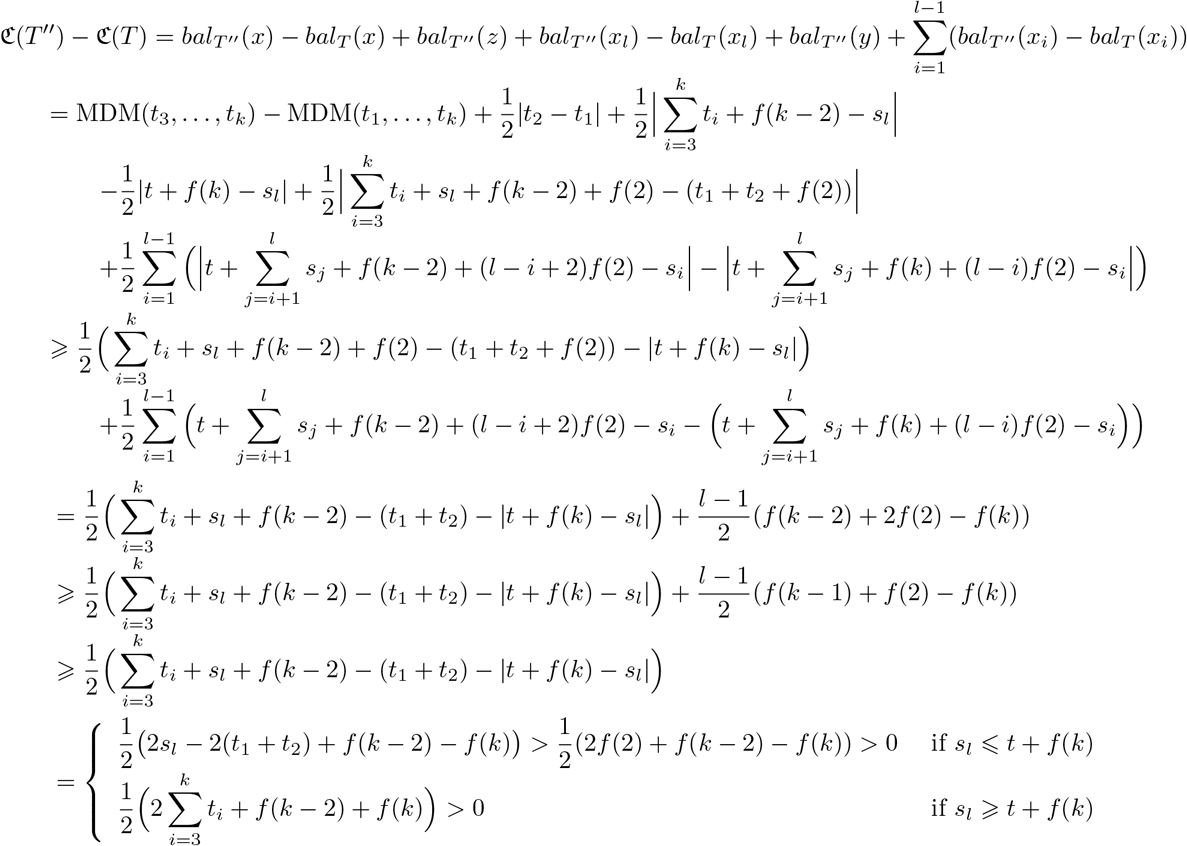

##### corollary 25.

*For every non-binary tree 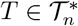, there always exists a binary tree 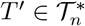 such that* ℭ(*T′*) *>* ℭ(*T*).

*Proof.* We shall prove by complete induction on the sum *S* of the degrees of the non-binary internal nodes in a tree 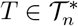 that there always exists a binary tree 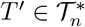 such that ℭ(*T′*) ⩾ ℭ(*T*). Moreover, it will be clear from the proof that if *T* isn’t binary, then *T′* can be chosen so that this inequality is strict.

The assertion to be proved by induction is obviously true if *S* = 0 (which means that *T* is binary), so assume that *S >* 0. Let *x* be an internal non-binary node of *T* such that all nodes in the path from the root *r* to *x*, except *x* itself, are binary.

If *x* is the root, then we apply Lemma 22 and we obtain a tree *T*_0_ with smaller *S* and larger ℭ. Then, by induction, there exists a binary tree *T′* such that ℭ(*T*) *<* ℭ(*T*_0_) ⩽ ℭ (*T″*).

If *x* is not the root *r*, let *r*, *x*_2_,…, *x_l_*, *x* be the path from the root to *x*: all these nodes except *x* are binary. By Lemma 23, if we rearrange the subtrees rooted at the children of the nodes *r*, *x*_2_,…, *x_l_* in increasing order of their *δ*-sizes, the ℭ value of the resulting tree increases without modifying the value of *S*; let *T̂* be the tree obtained in this way. Next, by Lemma 24, in *T̂* we can either prune a subtree *T*_1_ rooted at one child of *x* and regraft it to an arc in the path from *r* to *x* (adding a new binary node to the tree), or we can prune two subtrees *T*_1_, *T*_2_ rooted at two children of *x* and regraft their star *T*_1_ ⋆ *T*_2_ to an arc in the path from *r* to *x* (adding two new binary nodes to the tree), in both cases in such a way that the resulting tree *T*_0_ has a larger ℭ and a smaller *S*. Then, by induction, there exists a binary tree *T′* such that

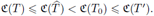

This finishes the proof by induction.

Therefore, the maximum ℭ value on 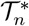 is reached at a binary tree, where, by Proposition 6 in the main text, it is equal to (*f* (0) + *f* (2))/2 times the colless index, with *f* (0) + *f* (2) ⩾ *f* (2) *>* 0. Then, since, by Lemma 1 in the main text, the maximum colless index of a binary tree with *n* leaves is reached exactly at the comb *K_n_*, the same is true for ℭ. So, the maximum value of ℭ on 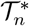 is reached exactly at *K_n_*, and it is

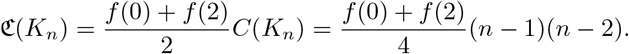

#### 0.2 Proof of the thesis of Theorem 18 for ℭ_***sd*,*f***_

The proof of Theorem 18 for *D* = *sd*, the sample standard deviation, is very similar to the one provided for *D* = MDM in the previous section, but simpler, because Lemmas 20 and 21 are replaced by Lemma 26 below, which guarantees that it is always enough to remove a suitable element in a non-constant numeric vector of length at least 3, in order to increase its variance. To simplify the notations, we shall denote in this section *δ_f_* and ℭ_*sd*,*f*_ by *δ* and ℭ, respectively, and we shall denote *bal_sd_*,*_f_* on a tree *T* by *bal_T_* or simply by *bal* when it is not necessary to specify the tree.

##### Lemma 26.

*For every* (*x*_1_,…, *x_n_*) ∈ ℝ *^n^ with n* ⩾ 3*, if x_i_ is the value in the set* {*x*_1_,…, *x_n_} closest to its mean, then*

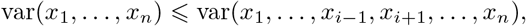

*and thus, taking positive square roots*,

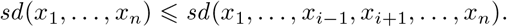

*Moreover, these inequalities are strict unless either x*_1_ =…= *x_n_ or n is even and* {*x*_1_,…, *x_n_*} *consists of n*/2 *copies of two different elements.*

*Proof.* Let *x̄* = (*x*_1_ +*…*+ *x_n_*)/*n* and, after rearranging *x*_1_,*…*, *x_n_* if necessary, assume that (*x_n_ – x̄*)^2^ ⩽ (*x_i_ – x̄*)^2^, for every *i* = 1,*…*, *n −* 1. We shall prove that

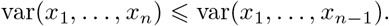

Indeed, let *x̄′* = (*x*_1_ + *…* + *x_n−_*_1_)/(*n −* 1). Then

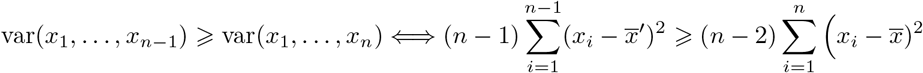

Now

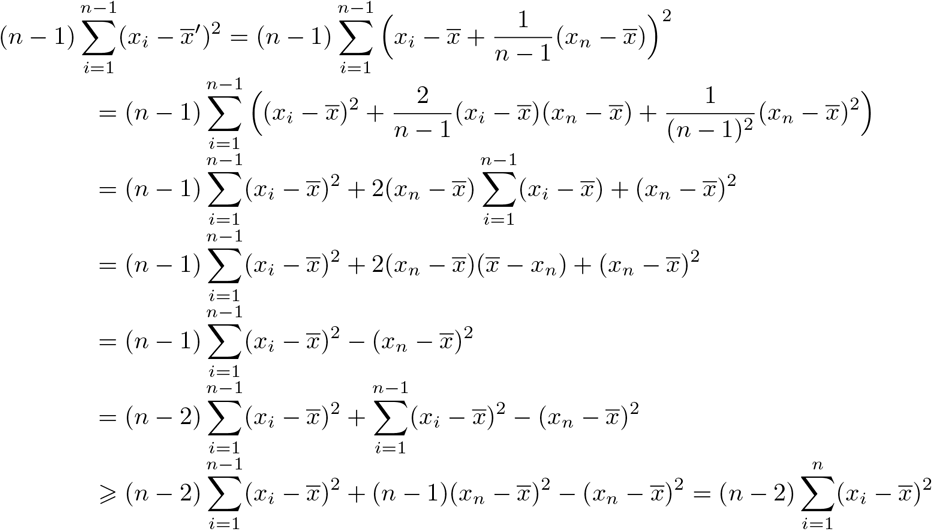

as we wanted to prove. This inequality is an equality if, and only if, (*x_i_* – *x̄*)^2^ = (*x_n_* – *x̄*)^2^ for every *i* = 1,…, *n −* 1, and it is easy to check that this condition holds exactly when either {*x*_1_ =…= *x_n_*} or *n* is even and {*x*_1_, *…*, *x_n_*} consists of *n/*2 copies of two different elements.

We prove now a series of lemmas that play in this proof the same role as Lemmas 22 to 24 in the last section.

##### Lemma 27.

*Let f be a mapping* ℕ→ℝ_⩾0_ *such that f >*(*k*) 0*, for every k* ⩾ 2*, and let T be a tree of the form T*_1_ ⋆..⋆*T_k_, with k* ⩾ 3*. if δ*(*T*_1_) *is the value in the set* {*δ*(*T*_1_),…, *δ*(*T_k_*) *closest to its mean, then taking the tree T′* = *T*_1_ ⋆ (*T*_2_ ⋆…⋆ *T_k_*) *depicted in the left-hand side of Fig. 17, we obtain that* ℭ(*T′*) *>* ℭ(*T*).

*Proof.* Let *t_i_* = *δ*(*T_i_*), for every *i* = 1,…, *k*. The only nodes in *T* or *T′* with different *bal* value in both trees are the roots and the new node *υ* in *T′*. Therefore,

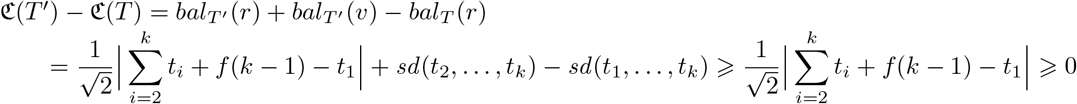

Now, the first inequality is strict unless either *t*_1_ = *t*_2_ =…= *t_k_* or (up to reordering the trees *T*_2_,…, *T_k_*) *k* = 2*m* ⩾ 4, *t*_1_ =…= *t_m_* and *t_m_*_+1_ =…= *t_k_*, and in both cases

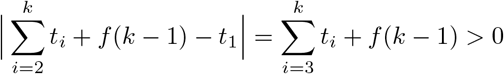

Therefore, we always have that ℭ(*T′*) *−* ℭ (*T*) *>* 0.

The proof of the following lemma is the same as that of Lemma 23 (up to replacing the fractions 1/2 by 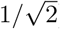), and we shall not repeat it here.

##### Lemma 28.

*Let f be a mapping* ℕ ⤍ℝ _⩾0_ *such that f* (2) *>* 0*. consider the trees T and T′ depicted in Fig. 18, where T′ is obtained from T by simply interchanging the subtrees T_l_ and T_l−_*_1_*. if δ*(*T_l_*) *< δ*(*T_l−_*_1_) *and δ*(*T_l_*) ⩽ *δ*(*T*_0_)*, then* ℭ(*T′*) *>* ℭ(*T*).

##### Lemma 29.

*Let f be a mapping* ℕ ⤍ℝ _⩾0_ *such that* 0 < *f* (*k*) < *f* (*k–*1) + *f* (2)*, for every k* ⩾ 3*, and let T be the tree depicted in Fig. 19, where l* ⩾ 1*, x*_1_ *is the root, all nodes in the path from x*_1_ *to x_l_ are binary, and k* ⩾ 3*. Assume moreover that δ*(*S*_1_) ⩽ *δ*(*S*_2_) ⩽… ⩽ *δ*(*S_l_*) *and that δ*(*T*_1_) *is the value in the set* {*δ*(*T*_1_),…, *δ*(*T_k_*)} *closest to its mean.*

(*a) if δ*(*S_l_*) ⩽ *δ*(*T*_1_)*, then the tree T′ depicted in Fig. 20, obtained by pruning the subtree T*_1_ *and regrafting it in the arc ending in x, is such that* ℭ(*T′*) *>* ℭ (*T*).
(*b) if δ*(*S_l_*) *> δ*(*T*_1_)*, then the tree T″ depicted in Fig. 20, obtained by pruning the subtree T*_1_ *and regrafting it in the arc ending in x_l_, is such that* ℭ(*T″*) *>* ℭ (*T′*).

*Proof.* For every *i* = 1,…, *l*, let *s_i_* = *δ*(*S_i_*) and, for every *i* = 1,…, *k*, *δ*(*T_i_*) = *t_i_*. Let, moreover, *t* = *t*_1_ +… + *t_k_*. So, we are assuming that *s*_1_ ⩽… ⩽ *s_l_* and that *sd* (*t*_1_,…, *t_k_*) ⩽ *sd* (*t*_2_,…, *t_k_*). Moreover, for every *i* = 1,…, *l*, we shall call *x_i_* the parent of the root of *S_i_* in all three trees *T*, *T′*, *T″*.

As far as assertion (a) goes, the only nodes in *T* or *T′* with different *bal* value in both trees are *x*_1_,…, *x_l_*, *x* and the new node *y*. Therefore:

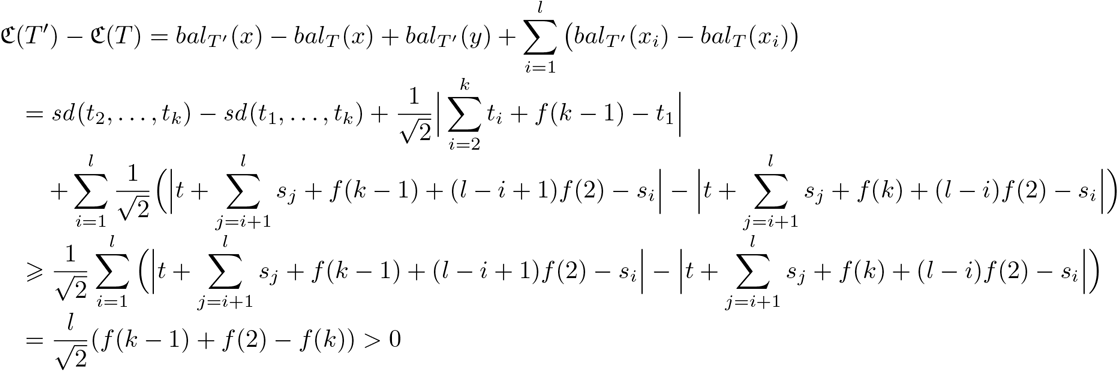

where, as in the proof of Lemma 24.(a1), the last equality is a consequence of the fact that, for every *i* = 1,…, *l*, *s_i_* ⩽ *s_l_* ⩽ *t*_1_ ⩽ *t*.

Let us prove now assertion (b). Again, the only nodes in *T* or *T″* with different *bal* value in both trees are *x*_1_,…, *x_l_*, *x* and the new node *y*. Therefore,

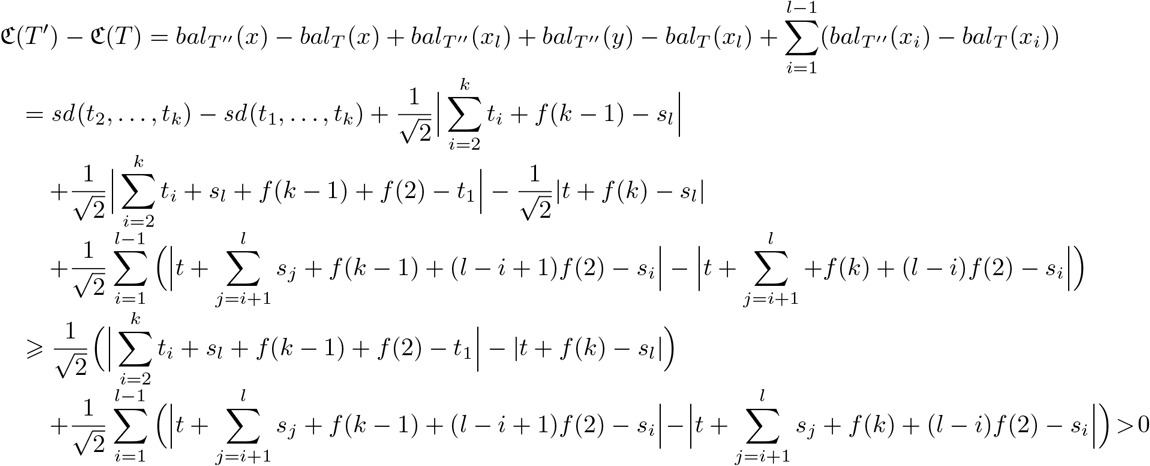

where the last strict inequality is derived as in the proof of Lemma 24.(a2).

Then, using Lemmas 27 to 29 and arguing as in the proof of corollary 25, we deduce that, for every non-binary tree 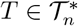, there always exists a binary tree such that ℭ(*T′*) *>* ℭ (*T*). Starting from this fact, the same argument that completes the proof of Theorem 18 for ℭ*_MDM,f_* also completes it for ℭ_*sd*,*f*_.

#### 0.3 Proof of the thesis of Theorem 18 for ℭ_var,*f*_

The proof is similar to those described in the previous two sections, using Lemma 26 and proving a series of lemmas that show how to increase the colless-like index of a non-binary tree by making it “more binary.” To simplify the notations, in this section, we shall denote *δ_f_* and ℭ_var,*f*_ by simply *δ* and ℭ, respectively, and we shall denote *bal_var,f_* on a tree *T* by *bal_T_* or simply by *bal* when it is not necessary to specify the tree.

##### Lemma 30.

*Let f be a mapping* ℕ *→* ℝ _⩾0_ *such that f* (*k*) *>* 0*, for every k* ⩾ 2*, and let T be a tree of the form T*_1_ ⋆… ⋆ *T_k_*, *with k* ⩾ 3*. if δ*(*T*_1_) *is the value in the set* {*δ*(*T*_1_),…, *δ*(*T_k_*)} *closest to its mean, then taking the tree T′* = *T*_1_ ⋆ (*T*_2_ ⋆… ⋆ *T_k_*) *depicted in the left-hand side of Fig. 17, we obtain that* ℭ(*T′*) *>* ℭ (*T*).

*Proof.* Let *t_i_* = *δ*(*T_i_*), for every *i* = 1,…, *k*. The only nodes in *T* or *T′* with different *bal* value in both trees are the roots and the new node *υ* in *T′*. Therefore,

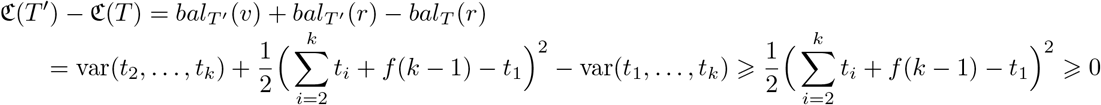

Now, the first inequality is strict unless either *t*_1_ = *t*_2_ =… = *t_k_* or (up to reordering the trees *T*_2_,…, *T_k_*) *k* = 2*m* ⩾ 4, *t*_1_ =… = *t_m_* and *t_m_*_+1_ =… = *t_k_*, and in both cases the last inequality is strict (cf. the proof of Lemma 27).

##### Lemma 31.

*Let f be a mapping* N ⤍ ℝ _⩾0_ *such that f* (2) *>* 0*. consider the trees T and T′ depicted in Fig. 18, where, in both trees, the nodes in the path connecting the root r with x are binary, and T′ is obtained from T by simply interchanging the subtrees T_l_ and T_l−_*_1_*. if δ*(*T_l_*) *< δ*(*T_l−_*_1_)*, then* ℭ(*T′*) *>* ℭ(*T*).

*Proof.* Let *t_i_* = *δ*(*T_i_*), for every *i* = 0,…, *l*, so that *t_l_* < *t_l−_*_1_. The only nodes in *T* or *T′* with different *bal* value in both trees are *x* and *y*, and therefore,

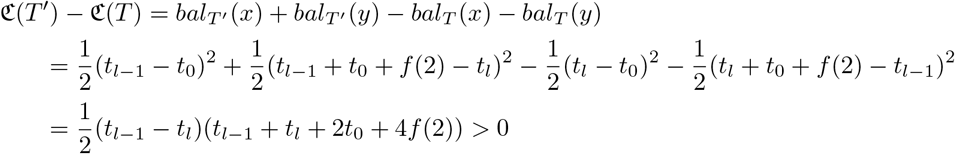

##### Lemma 32.

*Let f be a mapping* ℕ ⤍ℝ _⩾0_ *such that* 0 < *f* (*k*) < *f* (*k–*1) + *f* (2)*, for every k* ⩾ 3*, and let T be the tree depicted in Fig. 19, where l* ⩾ 1*, x*_1_ *is the root, all nodes in the path from x*_1_ *to x_l_ are binary, and k* ⩾ 3*. Assume moreover that δ*(*S*_1_) ⩽ *δ*(*S*_2_) ⩽… ⩽ *δ*(*S_l_*) *and that δ*(*T*_1_) *is the value in the set* {*δ*(*T*_1_),…, *δ*(*T_k_*)} *closest to its mean. Then:*

(*a) if δ*(*S_l_*) ⩽ *δ*(*T*_1_)*, then the tree T′ depicted in Fig. 20, obtained by pruning the subtree T*_1_ *and regrafting it in the arc ending in x, is such that* ℭ(*T′*) *>* ℭ (*T″*).

(*b) if δ*(*S_l_*) > *δ*(*T*_1_), *then the tree T″ depicted in Fig. 20, obtained by pruning the subtree T*_1_ *and regrafting it in the arc ending in x_l_, is such that* ℭ(*T″*) *>* ⩽ (*T*).

*Proof.* For every *i* = 1,…, *l*, let *s_i_* = *δ*(*S_i_*) and, for every *i* = 1,…, *k*, *δ*(*T_i_*) = *t_i_*, and let *t* = *t*_1_ +… + *t_k_*. We are assuming that *s*_1_ ⩽… ⩽ *s_l_* and that var(*t*_1_,…, *t_k_*) ⩽ var(*t*_2_,…, *t_k_*). Moreover, for every *i* = 1,…, *l*, we shall call *x_i_* the parent of the root of *S_i_* in all three trees *T*, *T′*, *T″*.

As far as assertion (a) goes, the only nodes in *T* or *T′* with different *bal* value in both trees are *x*_1_,…, *x_l_*, *x* and the new node *y*. Therefore,

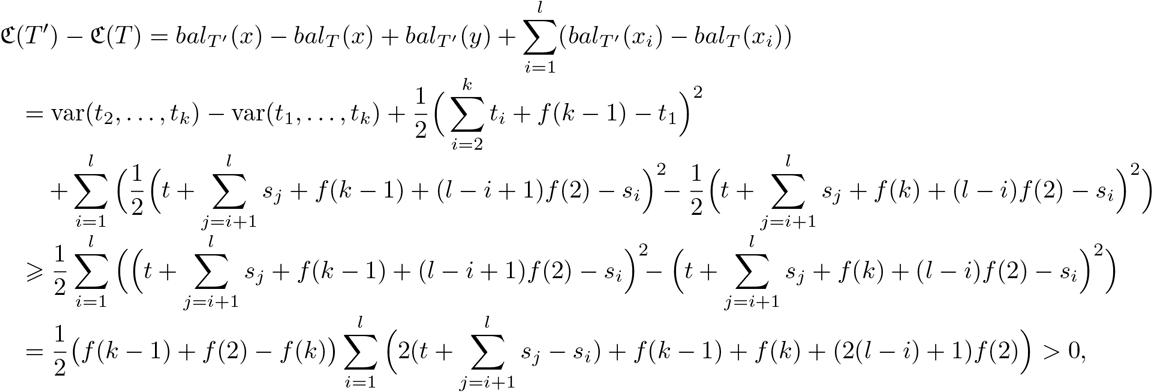

where this last expression is *>* 0 because *f* (*k–*1) + *f* (2)–*f*(*k*) *>* 0 and, for every *i* = 1,…, *l*, *s_i_* ⩽ *s_l_* ⩽ *t*_1_ ⩽ *t*.

Let us prove now assertion (b). Again, the only nodes in *T* or *T″* with different *bal* value in both trees are *x*_1_,…, *x_l_*, *x* and the new node *y*. Therefore,

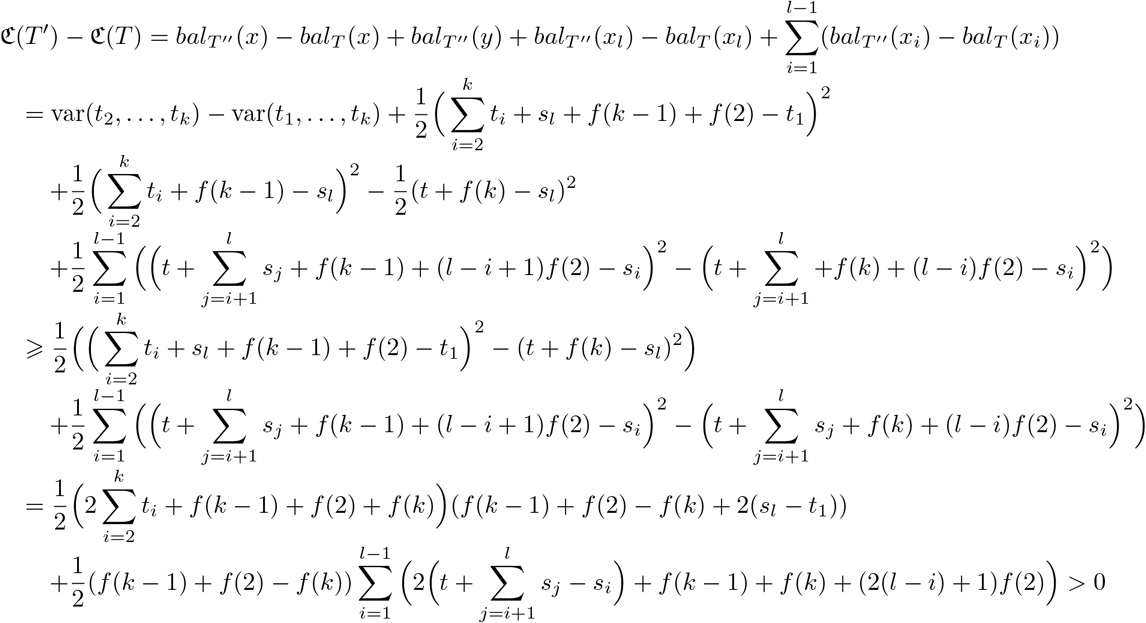

where this last expression is *>* 0 because *f* (*k −* 1) + *f* (2) *− f* (*k*) *>* 0, *s_l_ > t*_1_ and, for every *i* = 1,…, *l −* 1, *s_i_* ⩽ *s_l_*.

Then, using Lemmas 30 to 32 and arguing as in the proof of corollary 25, it can be proved that, for every non-binary tree 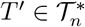, there always exists a binary tree 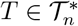 such that ℭ (*T′*) *>* ℭ (*T*). Therefore, the maximum ℭ value is reached at some binary tree. Since, for binary trees *T*, 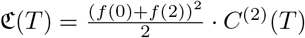 (see Proposition 7 in the main text), and *f* (0) + *f* (2) *>* 0, it remains to prove that the binary tree in 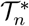 with maximum *c*^(2)^ is exactly the comb. The proof of this fact follows closely that of Lemma 1 in the main text.

##### corollary 33.

*For every binary tree 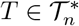, then c*^(2)^(*K_n_*) *> c*^(2)^(*T*).

*Proof.* Using the argument of the proof of Lemma 1 in the main text, it is enough to prove that if *T* and *T′* are the trees depicted in Fig. 6 in the main text, then, under the assumptions therein, *c*^(2)^(*T^′^*) *> c*^(2)^(*T*). And, indeed (using the notations therein),

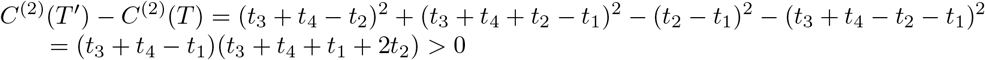

where the last inequality holds because, by assumption, *t*_1_, *t*_2_, *t*_3_, *t*_4_ > 0 and *t*_1_ + *t*_2_ ⩽ *t*_3_ + *t*_4_.

Now, it is straightforward to check that

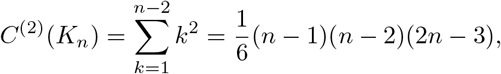

from where we obtain

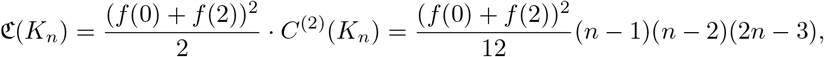

as we claimed in the statement.

#### 0.4 Proof of the thesis of Theorem 19 for 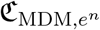

To simplify the notations, we shall denote in this section 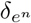 and 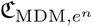 by *δ* and ℭ, respectively, and we shall denote *bal_MDM,f_* on a tree *T* by *bal_T_* or simply by *bal* when it is not necessary to specify the tree.

##### Lemma 34.

*Let n_1_*,…,*n_k_* (ℕ>0 *and n* = *n*_1_ +… + *n_k_*, *and assume that* 2 ⩽ *k* ⩽ *n −* 1*. Then*

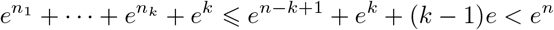

*Moreover, the first inequality is strict unless there is at most one exponent n_i_* ⩾ 2.

*Proof.* To begin with, notice that if 1 ⩽ *x* ⩽ min(*a*, *b*), then

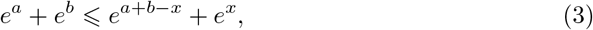

because

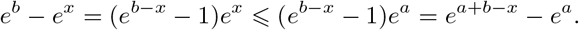

Moreover, if *x <* min(*a*, *b*) then the inequality is clearly strict.

Now, applying *k −* 1 times inequality (3) with *x* = 1, we obtain

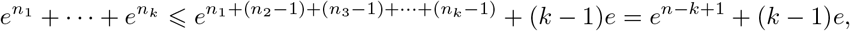

which implies the first inequality. Moreover, this inequality is strict unless all *n_i_* but, at most, one are 1, because (3) is strict if *x < a* and *x < b*, which in this situation is translated to the existence of at least two *n_i_, n_j_* greater than 1.

As far as the second inequality goes, since 2 ⩽ *k* ⩽ *n −* 1, and hence, in particular, *n* ⩾ 3 (and using, in the second inequality, that *x* + 1 *< e^x^* for every *x ∈* R_*>*0_), we have

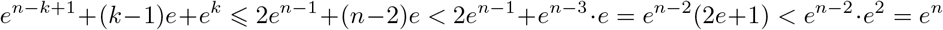

*e^n−k^*^+1^ +(*k−*1)*e*+*e^k^* ⩽ 2*e^n−^*^1^ +(*n−*2)*e <* 2*e^n−^*^1^ +*e^n−^*^3^ *⋅e* = *e^n−^*^2^(2*e*+1) *< e^n−^*^2^ *⋅e*^2^ = *e^n^* as we claimed.

##### Lemma 35.

*Let n*_1_,…, *n_k_*, *l ∈* ℕ *be such that k* ⩾ 1, *k* + *l* ⩾ 2, *each n_i_* ⩾ 2, *and let n* = *n*_1_…+ *n_k_. Then*

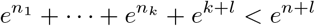

*Proof.* The case *l* = 0 is a particular instance of the last lemma. So, we assume henceforth that *l* ⩾ 1. if *k* = 1, so that *n* = *n*_1_, then applying inequality (3) with *x* = 2 we have

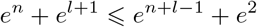

and the right hand side term is smaller than *e^n^*^+*l*^ because *n* + *l* ⩾ 3.

Assume finally that *k* ⩾ 2. Applying *k −* 1 times inequality (3) with *x* = 2, we obtain

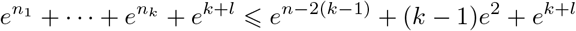

Now, since 2 ⩽ *k* ⩽ *n/*2, and hence *n* ⩾ 4,

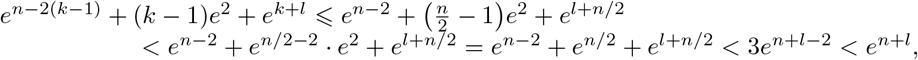

as we claimed.

##### Lemma 36.

*The largest e^n^-size of a tree in 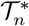, and it is reached exactly at the star FS_n_.*

*Proof.* The cases *n* = 1, 2 are obvious, because 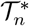 consists of a single tree. Let now *n* ⩾ 3. We shall prove that for every 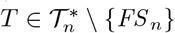, there is a tree 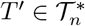 with *δ*(*T′*) *> δ*(*T*). This shows that no tree other than *FS_n_* can have the maximum *e^n^*-size.

So, let 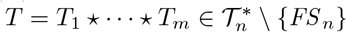, with *m* ⩾ 2. Let *l* ⩾ 0 be such that, for every *i* = 1,…, *l*, the subtree *T_i_* consists of a single node, and, for every *i* = *l* + 1,…, *m*, *T_i_* = *T_i,_*_1_ ⋆… ⋆ *T_i_*,*_n_i* with *n_i_* ⩾ 2; cf. Fig. 22. Since *T ≠ FS_n_*, *l* < *m.* Let now *T′* be the tree

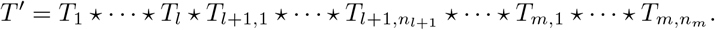

Then,

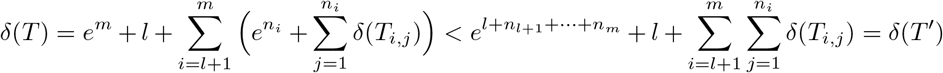

by Lemma 35.

##### Lemma 37.

*For every*
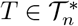 *with n*≠ 1, 3,

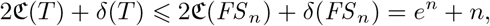

*and the inequality is strict if T ≠ FS_n_.*

*When n* = 1*, the maximum of* 2ℭ + *δ is* 2c(*FS*_1_) + *δ*(*FS*_1_) = 1, *and when n* = 3, *this maximum is* 2ℭ (*K*_3_) + *δ*(*K*_3_) = 3*e*^2^ + 4 *and it is reached exactly at K*_3_.

*Proof.* The cases *n* = 1, 2 are obvious, because then 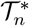 consists of a single tree, and the cases *n* = 3, 4, 5 can be checked in the tables in supplementary file S2. Notice that 1 < *e*^1^ + 1 and 3*e*^2^ + 4 *<* (*e*^3^ + 3) + 4; we shall use these two inequalities below. We prove now the general case *n* ⩾ 6 using the cases *n* = 1,…, 5 and complete induction on *n*.

Let *T* = *T*_1_ ⋆…⋆ *T_k_*, with *k* ⩾ 2 and 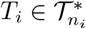 for every *i* = 1,…, *k*, so that *n* = *n*_1_ +…+ *n_k_* ⩾ 6. if *k* = *n*, then *n_i_* = 1 for every *i* and *T* = *FS _n_*, in which case 2ℭ(*T*) + *δ*(*T*) = *e^n^* + *n*. So, we shall assume that *k* ⩽ *n*–1, and we shall prove that, in this case 2ℭ (*T*) + *δ*(*T*) < *e^n^* + *n*

After renumbering the subtrees *T_i_* if necessary, assume that there exists *l* ⩾ 0 such that *n_i_* = 3 for every *i* ⩽ *l* and *n_i_* ≠ 3 for every *i* = *l* + 1,…, *k*. We shall prove first of all that

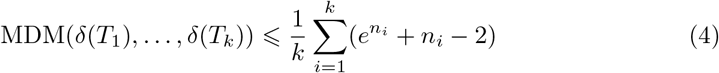

Indeed, let *M* = *Median*(*δ*(*T*_1_),…, *δ*(*T_k_*)). Then,

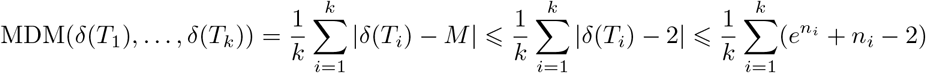

where the first inequality is due to the fact that *M* is the real number that minimizes the function 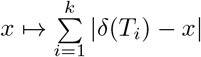, and the second inequality holds because 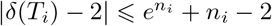 for every *i* = 1,…, *k*; on its turn, this inequality is a consequence, when *n_i_* ⩾ 2, of Lemma 36, and, when *n_i_* = 1, of the fact that if 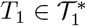, then *δ*(*T_i_*) = 1 and hence *|δ*(*T_i_*) *−* 2*|* = 1 *< e*^1^ + 1 *−* 2. Now,

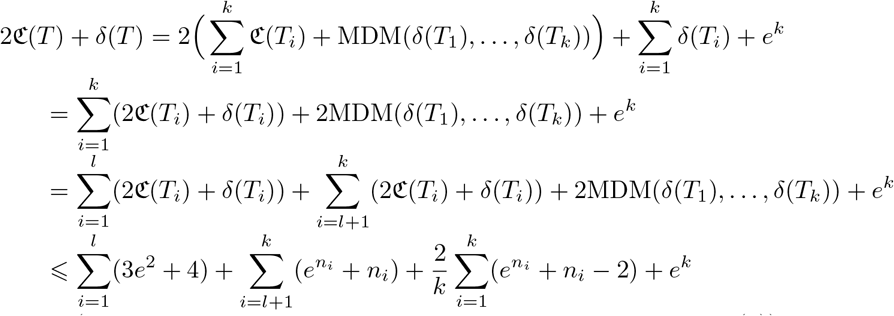

(by the case *n* = 3, the induction hypothesis, and inequality (4))

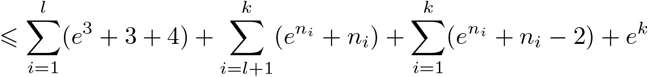

(because *k* ⩾ 2 and 3*e*^2^ + 4 *< e*^3^ + 3 + 4)

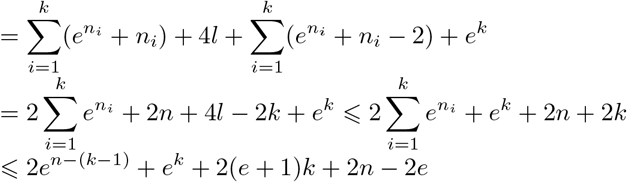

(by the first inequality in Lemma 34)

Thus, it remains to prove that, for every *n* ⩾ 6 and for every 2 ⩽ *k* ⩽ *n* − 1,

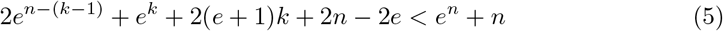

Since, for every *n* ⩾ 6, the function

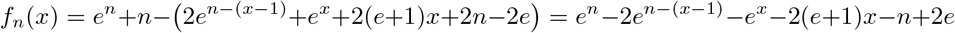

is concave, because 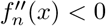, its minimum value on the closed interval [2, *n–*1] is reached at one of its ends. So, in order to prove inequality (5) for every *n* ⩾ 6 and for every *k* = 2,…, *n*–1, it is enough to prove that *f_n_*(2) *>* 0 and *f_n_*(*n*–1) *>* 0 for every *n* ⩾ 6. And, indeed

- *f_n_*(2) = *e^n^* − 2*e^n^*^−1^ − *n* − *e*^2^ − 2*e* − 4 > 0 because the function *g*(*x*) = *e^x^* − 2*e^x^*^−1^ − *x* − *e*^2^ − 2*e* − 4 is increasing on ℝ ⩾_2_ and *g*(5) *>* 0.
- *f_n_*(*n −* 1) = *e^n^ − e^n−^*^1^ *−* (2*e* + 3)*n −* 2*e*^2^ + 4*e* + 2 *>* 0 by a similar reason.

This finishes the proof of the statement.

Now we can proceed with the proof of Theorem 19.(a) for *D* = MDM. The cases *n* = 2, 3, 4, 5 can be checked in the tables in the supplementary file S2. Notice in particular that, when *n* = 4, the maximum is

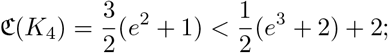

we shall use this inequality below. We prove now, using the cases *n* = 1,…, 5 and complete induction on *n*, that, for every *n* ⩾ 6,

*The tree in 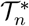 with maximum ℭ is FS*_1_ ⋆ *FS_n−_*_1_, *with*

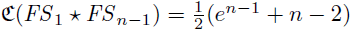

Recall that, as in the previous sections, Lemma 22 implies that the maximum ℭ value on 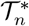 is reached at a tree with binary root. So, let 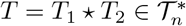, with 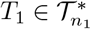 and 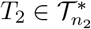. We must distinguish two cases:

*a*) Assume that *n*_1_ = 1, and therefore *n*_2_ = *n −* 1 ⩾ 5. in this case,

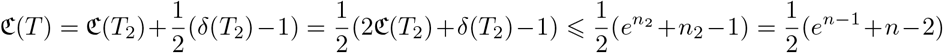

by Lemma 37. Moreover, the equality holds only when *T*_2_ = *FS _n−_*_1_.
*b*) Assume that *n*_1_, *n*_2_ ⩾ 2 and, without any loss of generality, that *δ*(*T*_2_) ⩽ *δ*(*T*_1_). Then,

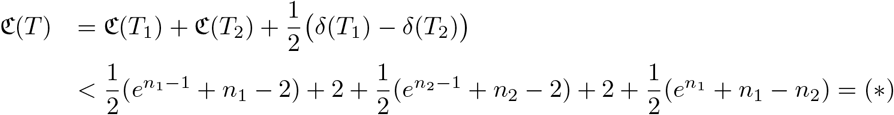

This inequality is due to the following facts. On the one hand, *n*_2_ ⩽ *δ*(*T*_2_) and 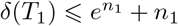, by Lemma 36, and hence 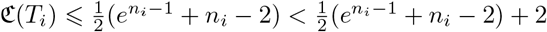. On the other hand, by the induction hypothesis, 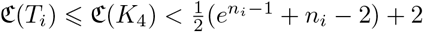, unless *n_i_* = 4, in which case we still have 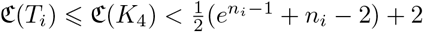.

Let us continue

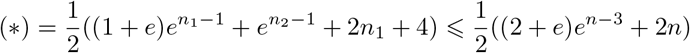

because *n*_1_, *n*_2_ ⩽ *n −* 2. So, it remains to prove that, for every *n* ⩾ 6,

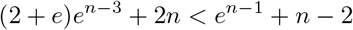

This is equivalent to

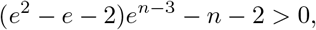

which is easy to prove, for instance noticing that *f* (*x*) = (*e*^2^ *− e −* 2)*e^x−^*^3^ *− x −* 2 is increasing on ℝ _⩾3_ and that *f* (5) *>* 0. This finishes the proof of Theorem 19 for *D* = MDM.

#### 0.5 Proof of the thesis of Theorem 19 for 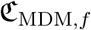

The proof of this case follows closely that of the case when *D* = MDM given in the last section. To begin with, it turns out that a key lemma similar to Lemma 37 also holds when *D* = *sd*. To simplify the notations, we shall denote in this section *δ_e^n^_* and 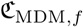 by *δ* and ℭ, respectively, and we shall denote *bal_sd_*_,*f*_ on a tree *T* by *bal_T_* or simply by *bal* when it is not necessary to specify the tree.

##### Lemma 38.

*For every 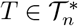 with n* ≠ 1, 3,

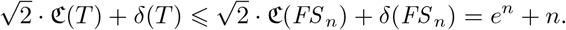

*and the inequality is strict if T ≠ FS_n_*.

*When n* = 1, *the maximum of* 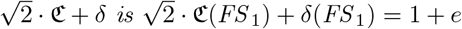, *and when n* = 3, *this maximum is* 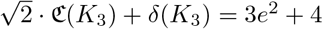.

*Proof.* The cases *n* = 1, 2 are obvious, and the cases *n* = 3, 4, 5 can be checked in Table 2 in the supplementary file S2. We shall use that 1 *< e*^1^ + 1 and the following inequalities:

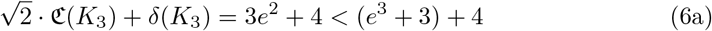

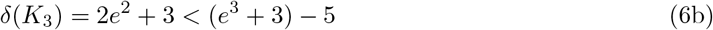

We prove the general case *n* ⩾ 6 by induction on *n* using an argument very similar to the one given in the proof of Lemma 37. Let *T* = *T*_1_ ⋆… ⋆ *T_k_*, with *k* ⩾ 2 and 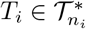 for every *i* = 1,…*, k*, so that *n* = *n*_1_ +… + *n_k_*. if *k* = *n*, then *n_i_* = 1 for every *i* and *T* = *FS_n_*, in which case 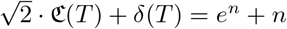. So, we shall assume that *k* ⩽ *n–*1. Without any loss of generality, we assume that there exists *l* ⩾ 0 such that *T_i_* = *K*_3_ if, and only if, *i* ⩽ *l*.

Now, it turns out that

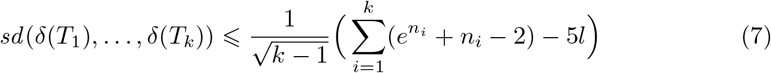

Indeed, let *m* = (*δ*(*T*_1_) +…+ *δ*(*T_k_*))*/k*. Then,

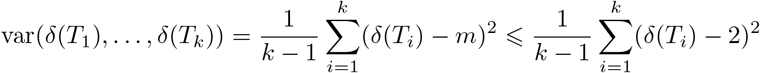

because *m* is the real number that minimizes the function 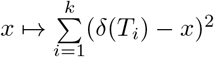. Taking

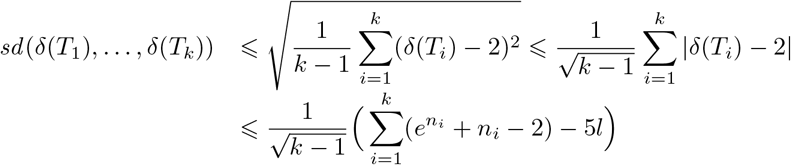

where the second last inequality is a consequence of Lemma 36, inequality (6b), and the fact, already used in the previous section, that if 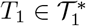, then |*δ*(*T*_1_)–2| = *e*–1 = *e*+1–2. Then

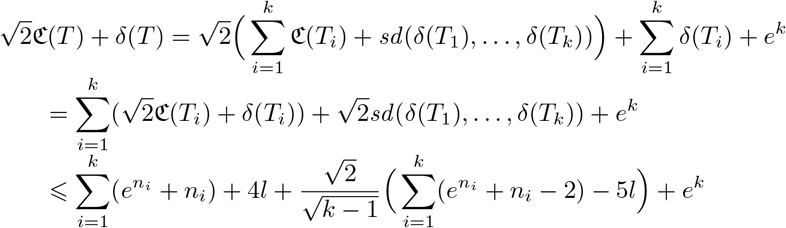

(by the induction hypothesis and inequalities (6a) and (7))

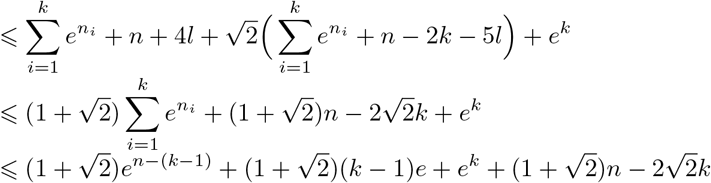

(because of the first inequality in Lemma 34)

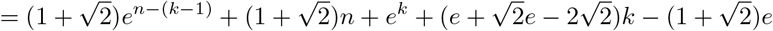

Thus, it remains to prove that, for every *n* ⩽ 6 and for every 2 ⩽ *k* ⩽ *n −* 1,

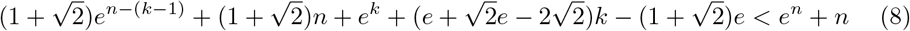

Now, for every *n* ⩾ 1, the function

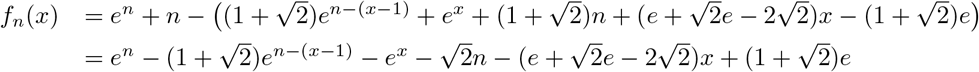

is concave, and therefore the minimum value of *f_n_*(*x*) on the closed interval [2, *n* − 1] will be reached at one of its ends. So, in order to prove inequality (8) for every *n* ⩾ 6 and every *k* = 2,…, *n* − 1, it is enough to prove that *f_n_*(2) > 0 and *f_n_*(*n* − 1) > 0 for every *n* ⩾ 6. And, indeed

- 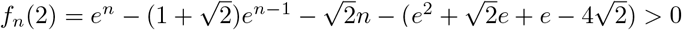 because the function 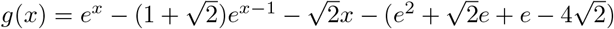 is increasing on ℭ _⩾ 3_ and *g*(5) *>* 0.
- 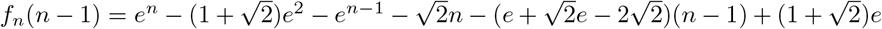 by a similar reason.

This finishes the proof of the lemma.

From here on, the proof of Theorem 19 for *D* = *sd* proceeds as the one for *D* = MDM given in the previous section, using Lemma 38 instead of Lemma 37; to ease the task of the reader we provide it. The cases *n* = 2, 3, 4, 5 can be checked in Table 2 in the supplementary file S2. Notice in particular that, when *n* = 4, the maximum ℭ value is

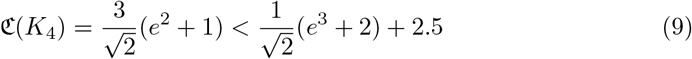

we shall use it below.

We prove now, using the cases *n* = 1,…, 5 and complete induction on *n*, that, for every *n* ⩾ 6,

*The tree in 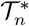 with maximum* ℭ *is FS*_1_ ⋆ *FS _n−_*_1_, *with*

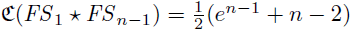

To begin with, notice that Lemma 27 implies that the maximum ℭ value on 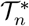 is reached at a tree with binary root. So, let 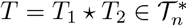, with 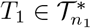 and 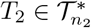. We must distinguish two cases:

a) Assume that *n*_1_ = 1, and therefore *n*_2_ = *n −* 1 ⩾ 5. in this case,

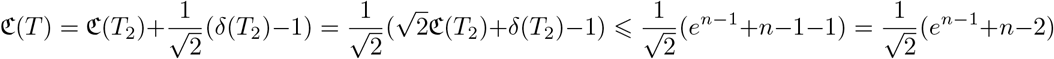

by Lemma 37. Moreover, the equality holds only when *T*_2_ = *FS*_*n* − 1_.
b) Assume that *n*_1_, *n*_2_ ⩾ 2 and, without any loss of generality, that *δ*(*T*_2_) ⩽ *δ*(*T*_1_). Then,

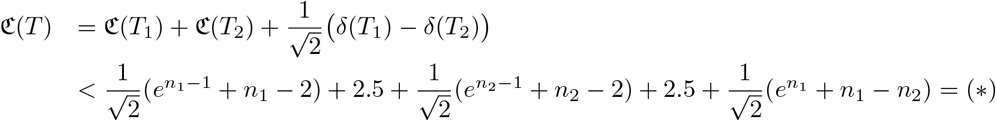

This inequality is due to the following facts. On the one hand, *n*_2_ ⩽ *δ*(*T*_2_) and 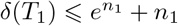, by Lemma 36, and hence 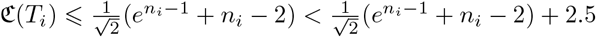. On the other hand, by the induction hypothesis, 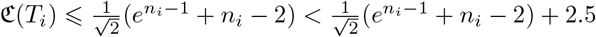, unless *n_i_* = 4, in which case we still have 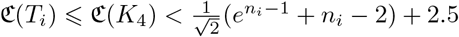 by inequality (9).

Let us continue

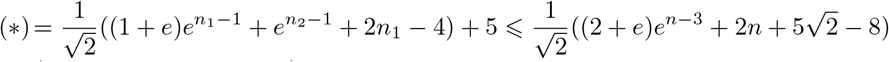

(because *n*_1_, *n*_2_ ⩽ *n* − 2)

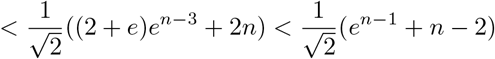

because (2 + *e*)*e*^*n* − 3^ + 2*n* < *e*^*n* − 1^ + *n* − 2, as it was proven in the last step of the proof of Theorem 19 for *D* = MDM in the last section. This finishes the proof of Theorem 19 for *D* = *sd*.

#### 0.6 Proof of the thesis of Theorem 19 for 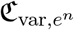

The stated maximum value of ℭ_var,*e*_*n* on 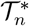, for *n* = 2,…, 5, can be checked in Table 2 in the supplementary file S2. As far as the case when *n* ⩾ 6, it is a direct consequence of the corresponding result for *D* = *sd*, established in the previous section.

Indeed, to begin with, notice that, since, for every node *υ* in a tree *T*, *bal*_var,*f*_ (*υ*) = *bal*_*sd,f*_ (*υ*)^2^, we have that, for every tree *T*,

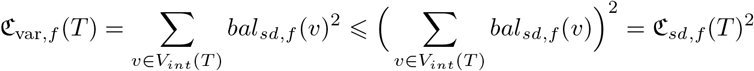

So, for every 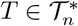 with *n* ⩾ 6,

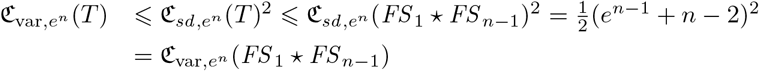

where the second inequality is strict if *T* ≠ *FS* _1_ ⋆ *FS*_*n* − 1_.

#### Supplementary file S2: Some tables

**Table 1.**
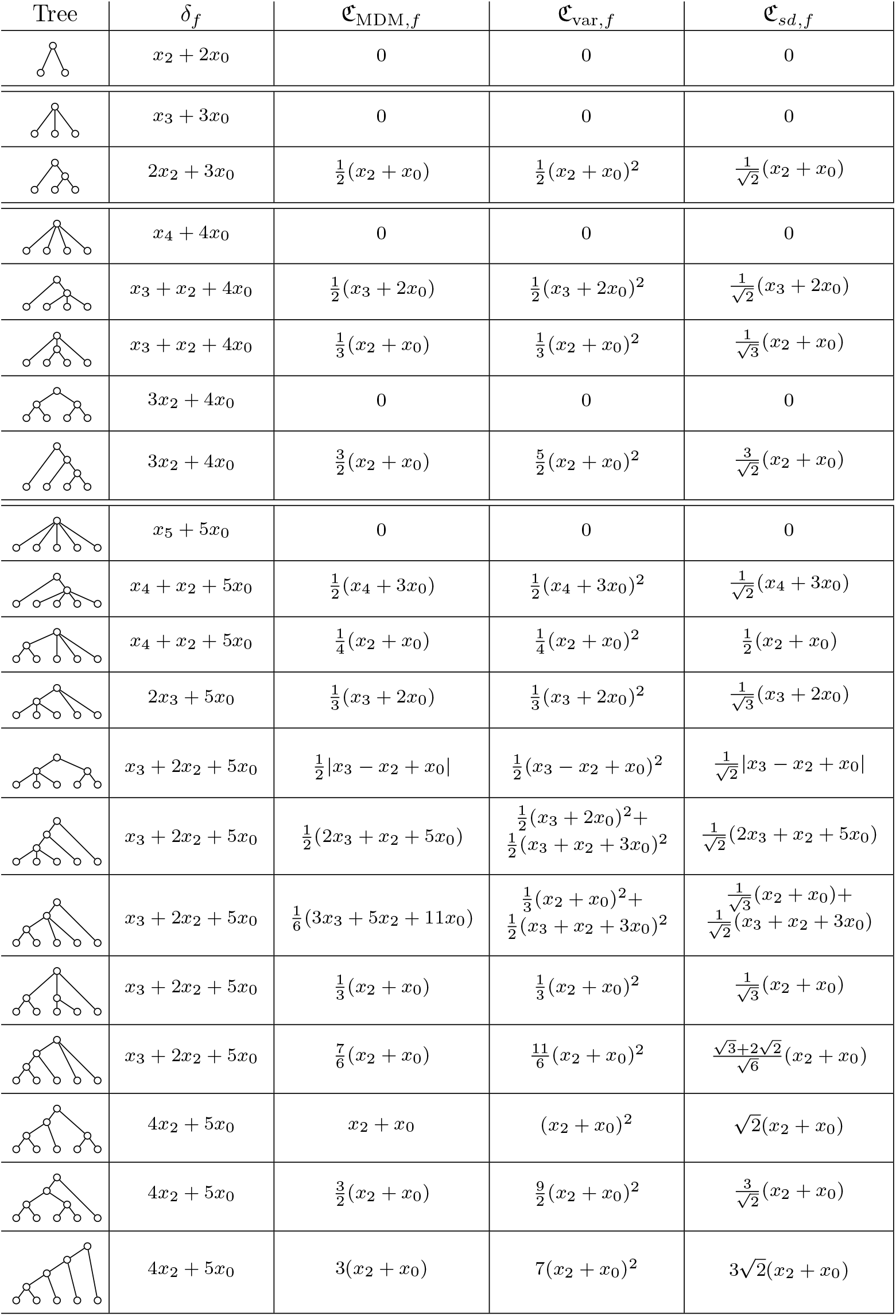
Abstract values of *δ*_*f*_, ℭ_MDM,*f*_, ℭ_var,*f*_, and ℭ_*sd,f*_ on 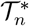 for *n* = 2, 3, 4, 5. We denote *f* (*i*) by *x_i_*.

**Table 2.** Numerical values (rounded to 4 decimal places) of *δ_f_* (for *f* (*n*) = ln(*n* + *e*) and *f* (*n*) = *e^n^*) and of ℭ_*D*,*f*_ (for every combination of *D* = MDM, var or *sd* and *f* (*n*) = ln(*n* + *e*) or *f* (*n*) = *e^n^*) on 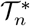, for every *n* = 2, 3, 4, 5. To win some horizontal space, in the subscripts we have replaced MDM by simply M, and we have denoted ln(*n* + *e*) by ln. The columns labelled “Pos.” give the position of the tree in its 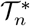 in increasing order of the colless-like balance index corresponding to the column on its left. The rows are sorted, for each *n*, in increasing order of ℭ_MDM,ln_.

| Tree | $\delta_{\ln}$ | $\mathfrak{C}_{M,\ln}$ | Pos. | $\mathfrak{C}_{\text{var},\ln}$ | Pos. | $\mathfrak{C}_{sd,\ln}$ | Pos. | $\delta_{e^n}$ | $\mathfrak{C}_{M,e^n}$ | Pos. | $\mathfrak{C}_{\text{var},e^n}$ | Pos. | $\mathfrak{C}_{sd,e^n}$ | Pos. |
| --- | --- | --- | --- | --- | --- | --- | --- | --- | --- | --- | --- | --- | --- | --- |
|  | 3.5514 | 0 | (1) | 0 | (1) | 0 | (1) | 9.3891 | 0 | (1) | 0 | (1) | 0 | (1) |
|  | 4.7437 | 0 | (1) | 0 | (1) | 0 | (1) | 23.0855 | 0 | (1) | 0 | (1) | 0 | (1) |
|  | 6.1029 | 1.2757 | (2) | 3.2549 | (2) | 1.8041 | (2) | 17.7781 | 4.1945 | (2) | 35.1881 | (2) | 5.9320 | (2) |
|  | 5.9048 | 0 | (1) | 0 | (1) | 0 | (1) | 58.5982 | 0 | (1) | 0 | (1) | 0 | (1) |
|  | 8.6543 | 0 | (1) | 0 | (1) | 0 | (1) | 26.1672 | 0 | (1) | 0 | (1) | 0 | (1) |
|  | 7.2951 | 0.8505 | (2) | 2.1700 | (2) | 1.4731 | (2) | 31.4746 | 2.7964 | (2) | 23.4588 | (2) | 4.8434 | (2) |
|  | 7.2951 | 1.8718 | (3) | 7.0075 | (3) | 2.6472 | (3) | 31.4746 | 11.0428 | (3) | 243.8855 | (4) | 15.6168 | (3) |
|  | 8.6543 | 3.8272 | (4) | 16.2747 | (4) | 5.4124 | (4) | 26.1672 | 12.5836 | (4) | 175.9407 | (3) | 17.7959 | (4) |
|  | 7.0436 | 0 | (1) | 0 | (1) | 0 | (1) | 153.4132 | 0 | (1) | 0 | (1) | 0 | (1) |
|  | 9.8466 | 0.5961 | (2) | 0.7107 | (2) | 0.8430 | (2) | 39.8636 | 6.8482 | (4) | 93.7968 | (5) | 9.6849 | (4) |
|  | 8.4563 | 0.6379 | (3) | 1.6275 | (3) | 1.2757 | (3) | 66.9872 | 2.0973 | (2) | 17.5941 | (2) | 4.1945 | (2) |
|  | 9.8466 | 0.8505 | (4) | 2.1700 | (4) | 1.4731 | (4) | 39.8636 | 2.7964 | (3) | 23.4588 | (3) | 4.8434 | (3) |
|  | 8.4873 | 1.2479 | (5) | 4.6717 | (5) | 2.1614 | (5) | 45.1711 | 7.3618 | (5) | 162.5903 | (7) | 12.7511 | (6) |
|  | 8.4563 | 2.4524 | (6) | 12.0287 | (7) | 3.4682 | (6) | 66.9872 | 28.7991 | (12) | 1658.7734 | (12) | 40.7280 | (12) |
|  | 11.2058 | 2.5514 | (7) | 6.5099 | (6) | 3.6083 | (7) | 34.5562 | 8.3891 | (6) | 70.3763 | (4) | 11.8639 | (5) |
|  | 9.8466 | 2.9767 | (8) | 11.9348 | (8) | 4.7503 | (8) | 39.8636 | 9.7872 | (7) | 129.0231 | (6) | 15.6188 | (7) |
|  | 11.2058 | 3.8272 | (9) | 29.2944 | (11) | 5.4124 | (9) | 34.5562 | 12.5836 | (8) | 316.6932 | (8) | 17.7959 | (8) |
|  | 9.8466 | 3.9980 | (10) | 21.9842 | (9) | 5.9244 | (10) | 39.8636 | 18.0336 | (9) | 487.8092 | (9) | 26.3922 | (9) |
|  | 9.8466 | 5.0194 | (11) | 26.8218 | (10) | 7.0985 | (11) | 39.8636 | 26.2801 | (11) | 708.2359 | (11) | 37.1656 | (11) |
|  | 11.2058 | 7.6543 | (12) | 45.5691 | (12) | 10.8249 | (12) | 34.5562 | 25.1672 | (10) | 492.6338 | (10) | 35.5918 | (10) |

